# A Dynamic Bayesian Actor Model explains Endpoint Variability in Homing Tasks

**DOI:** 10.1101/2022.11.09.515854

**Authors:** Fabian Kessler, Julia Frankenstein, Constantin A. Rothkopf

## Abstract

Goal-directed navigation requires integrating information from a variety of internal and external spatial cues, representing them internally, planning, and executing motor actions sequentially. However, a comprehensive computational account of how these processes interact in an ambiguous, uncertain, and noisy environment giving rise to biases and variability observed in navigation behavior is currently unavailable. In this paper, we introduce an optimal control under uncertainty model, which provides a computational-level explanation of how landmarks and path integration interact and are combined to reduce variability in navigation. We apply our model to trajectory and end-point data from three previously published studies that employ a variant of the triangle completion task with landmarks. Contrary to observer models, which attribute human end-point variability to perceptual cue combination processes only, this dynamic Bayesian actor model provides a unifying account of a wide range of phenomena found in this task by considering variability in perception, action, and internal representations jointly. Taken together, these findings have wide-ranging implications for the analysis and interpretation of human navigation behavior, including resolution of seemingly contradictory results on cue integration in navigation.

## 1 Introduction

Navigation is one of the most fundamental goal-directed behaviors relevant to survival in animals and humans. As navigation is the cognitive ability to both plan and move from one place to a goal location [1], it involves dynamic interactions of multi-sensory perceptual, cognitive, as well as motor processes [2]. Human navigation performance has been shown to exhibit large variability [3–5], which has been associated with noise in sensory [6–8], representational [9, 10], and motor systems [11–13]. However, while previous research has investigated navigation behaviorally [3, 14, 15], its neuronal underpinnings [16, 17], prominently internal representations related to a cognitive map [18–20], and through computational modeling [21–24], a comprehensive account of uncertainty and variability involved in navigation is still elusive [25, 26]. Understanding how these systems interact and how navigational variability is caused by noise in these systems is crucial for understanding how the brain makes use of ambiguous, uncertain, and noisy information from different, sometimes conflicting, sources [27] to maintain a sense of location and direction while moving through the environment.

Information relevant to navigation may be collected from internal, e.g., motor cues or external sources, e.g., landmarks. Depending on cue availabilities, several navigational behaviors have been described, including path-integration [28], beaconing [29], landmark-based strategies [29–31], strategies based on route knowledge [32] or the use of a cognitive map [9, 18, 33]. Whether and how information from landmarks and path integration are combined when estimating the position or homing direction in navigation [34–41], is an ongoing debate. Paradigmatic experiments for studying these relationships involve variants of the triangle completion task [3], in which subjects navigate a triangular-shaped outbound path by picking up a sequence of three objects in a real [35, 37] or virtual environment [38–41] with landmarks. Subjects then have to return to the location of the first object (homing) under varying cue conditions, e.g., landmark cues only, path-integration cue only, or both cues simultaneously, leading to different patterns of trajectories and endpoint variability.

Current computational models of triangle completion tasks explain variability in homing responses as perceptual cue integration [35, 42] but offer no account as to how or why navigational variability arises in the first place. Under the assumption of optimal integration of cues, individual cues are combined according to their empirical relative reliabilities so that the integrated response is located between the responses to either cue alone but biased toward the more reliable cue. Further, the integration of variabilities observed in the two single cue conditions, i.e., landmark and self-motion, should yield lower endpoint variability in double cue conditions. While some studies found behavior in line with the predictions of Bayesian cue integration [35], others highlighted several discrepancies [37–39, 41]. As an example, for cue conflicts due to large landmark rotations, a single cue may determine the response direction, but response variability was reduced at the same time [38, 43], while for small landmark rotations, the response direction was biased toward the path-integration cue [39]. A recent study [44] suggested accounting for discrepancies to Bayesian cue integration by postulating a strong prior over target locations and additional behavioral costs but still framing the task as perceptual cue integration.

However, cue integration models are ideal observer models [45], i.e., they are purely perceptual and do not incorporate uncertainty in either motor actions [46] or internal models [27]. More importantly, they only consider a single percept or action, i.e., the selection of a single response location [35, 39] or direction [38]. But navigation is inherently a sequential visuomotor behavior, which unfolds over space and time, and requires the actor to continuously integrate sensory uncertainty, internal model uncertainty, motor variability, and behavioral costs to plan subsequent actions [26, 47–50]. Thus, variability in trajectories and endpoints results from multiple interacting uncertainties and variabilities that accrue over time, which the actor shapes actively through behavior. Here, we present a computational model of human spatial navigation based on optimal control under uncertainty [51, 52]. The model involves continuous states and actions, noise in sensory signals, internal representations and actions, non-linear egocentric motion and observation models, as well as a dynamic, internal, uncertain allocentric map. By assuming that subjects employ belief state planning [53], i.e., that they continuously plan their actions based on subjective, internal beliefs combining different, uncertain sources of information, this model predicts humans’ variability in trajectory endpoints observed in multiple studies [35, 38, 39]. The dynamic Bayesian actor model provides a unifying account of both state estimation (Where am I? Where is my goal?) as well as planning and control (Where should I go? How do I get there?) while taking uncertainty in perception, action, and internal representations into account. The results show that perception and action in spatial navigation must be considered jointly because they are inseparably intertwined [54, 55].

## 2 Results

### Task Description & Behavioral Data Analysis

To model behavior in homing tasks, we obtained the environmental layouts, methodological details, and behavioral data from three previously published studies [35, 38, 39] (see Methods; Figure S1 and Table 1). In all studies, human subjects performed a triangle completion task in either a real or a virtual environment containing landmarks (see Figure 1A). On each trial, subjects viewed the environment from their starting position. They then walked an outbound path through a sequence of three goal locations (see Figure 1B). Once they reached the last target location, they had to wait 8-20 seconds, during which the environment was invisible. When the environment reappeared, internal or external cues were manipulated under one of four cue conditions. Subjects had to return to the remembered first goal location leading to different patterns of endpoint variability (see Figure 1C). For the behavioral analysis, homing responses were are at the respective true target location (see Figure 1D). Based on the distributions of endpoints in each of the separate homing conditions, response variability is computed as the standard deviation of the euclidean distances (or heading directions) between the response locations for each trial to the mean response location (or heading), see Figure 1E. Response variability for the combined cue condition (see Figure 1E; top left; green vs. gray bar) is predicted from response variability in single cue conditions according to the perceptual cue integration model for euclidean distance [35, 39] or heading direction [38]. Mean responses in conflict conditions (see Figure 1E; top right) are used to predict relative response proximity to either landmark or self-motion defined target locations and compared with optimal predictions based on response variability (see Figure 1E; bottom).

**Table 1.** Task parameters were extracted from the methods section of the respective paper, the trajectory data (if available), or further clarified in correspondence with the studies original authors.

| Study / Parameter | Nardini 2008 | Chen 2017 | Zhao 2015 |
| --- | --- | --- | --- |
| Environment Size | 6 x 10 m | 6.9 x 8.8m | 10 x 10m |
| Viewing Modality | Regular vision | HMD / VR | HMD / VR |
| Field of view | 180° | 60° | 63° |
| Number of Landmarks | 3 | 1 or 3 | 3 (proximal vs. Distal) |
| Target visibility at start | all at once | sequentially | sequentially |
| Target Jitter | none | plus or minus 0.4m | none |
| Homing distance | 1.75m | 3.6m | 2.15m |
| Number of trials per condition (per subject) | 4 (16 total) | 10 (40 total) | 40 (360 total) |
| Waiting Time (all conditions) | 8 seconds | 20 seconds | 10 seconds |
| Landmark rotation (conflict) | 15° (left and right) | 15° (left and right) | 15°, 30°, 45°, 90° |
| Disorientation Turning Speed (landmark) | ~ 90°/s | ~ 90°/s | ~ 73°/s |
| Disorientation Position (landmark) | 0.5m | extracted from trajectory data | extracted from trajectory data |
| Reorientation prior homing (landmark) | yes | no | no |

**Figure 1.**
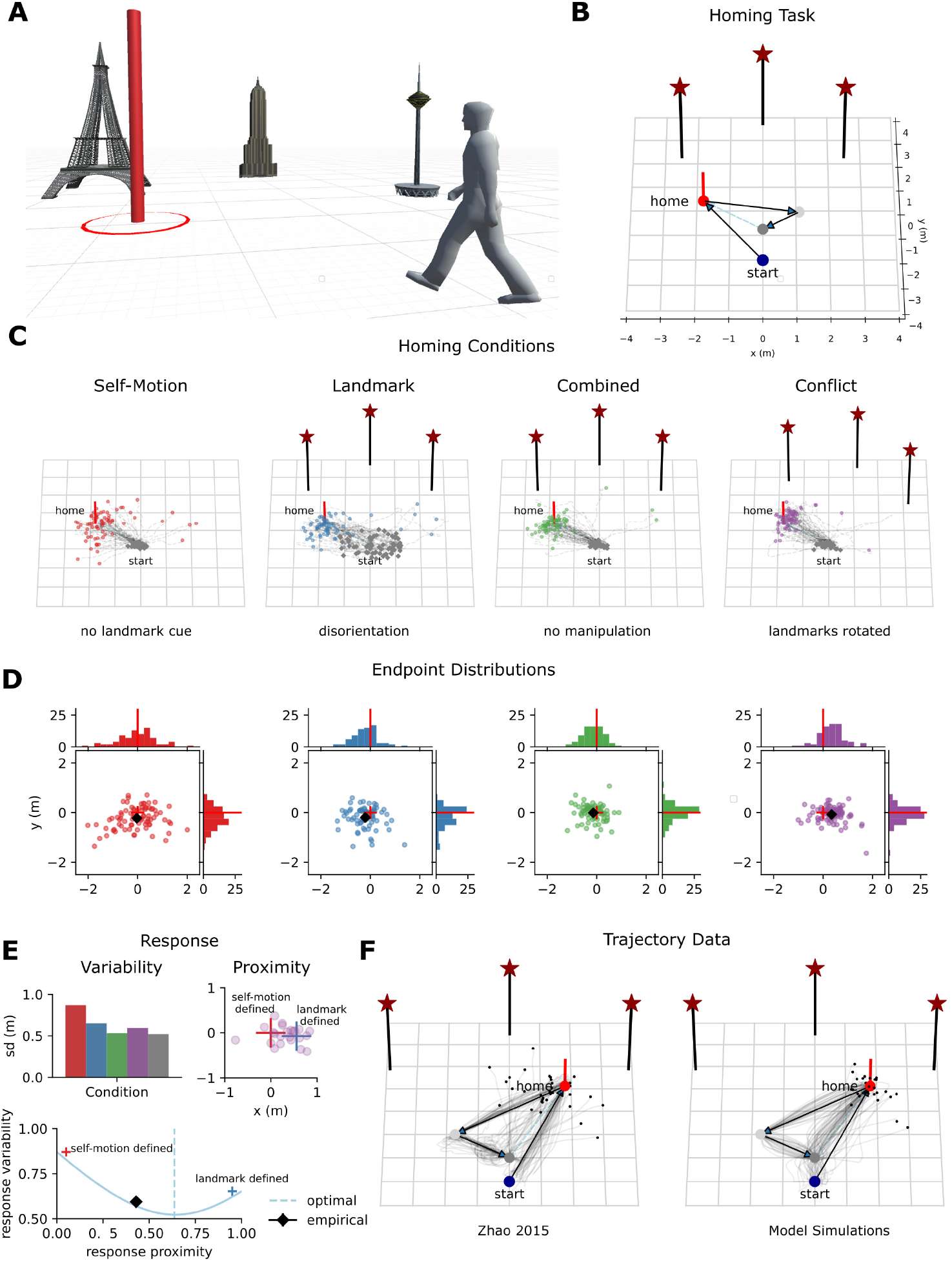
Homing task with landmarks, ideal observer account & computational model simulations. (A) Schematic of a typical homing experiment within a real [35] or virtual [38, 39] environment with landmarks. (B) Exemplary task environment for the triangle completion homing task with three landmarks. Subjects walk from a start position through a sequence of three goal locations before returning to the first location (home, shown in red). Walked routes vary on a trial-by-trial basis, whereas landmark locations remain stable across trials. (C) Experimental manipulation of internal and external cues in different task conditions prior to homing leads to different endpoint distributions. In the self-motion condition, subjects navigate without landmark cues (or any visual information). In the landmark condition, participants are disoriented by turning on a swivel chair to disrupt information about their prior heading direction and position (see blue dots). In the combined condition, all cues are available and undisrupted. In the conflict condition, landmarks are covertly rotated by 15° [35, 39] or up to 135° [38] around the location of the last target. Exemplary data from Chen 2017 [39]. (D) Subjects homing responses from different target locations are centered within a common coordinate frame, and outliers are removed. (E) Ideal Observer account of spatial navigation in the homing task [35]. Response variability is computed as the standard deviation of euclidean distances [35, 39] (or heading direction [38]) to mean response locations for each condition. Left: Response variability for different cue conditions. Variability in landmark (blue) and self-motion (red) conditions predict reduced variability in combined conditions (green) according to perceptual cue integration models [35] (gray). Right: In conflict conditions, landmarks are rotated around the third target location, creating a conflict between landmark and self-motion cues. Biased homing responses in conflict conditions are used to determine how much subjects relied on either of the two cues by computing relative response proximity to either cue. Bottom: If cues are combined optimally, response variability is reduced compared to single cue conditions, i.e., landmark and self-motion, and the optimal response location is biased toward the more reliable cue. Optimal predictions in terms of response variability and response proximity of the ideal observer model are compared to the behavior observed in the task. (F) Simulations from our dynamic Bayesian actor model compared to trajectory data of Zhao 2015 [38].

### Dynamic Bayesian Actor Model

The homing task described above encompasses a wide range of navigation scenarios, including walking towards visible and invisible targets while keeping track of one’s position and building up an internal representation of space, orienting based on land-marks in the environment, and finally walking back to the remembered home location. During navigation, the actual position of the subject and the position of landmarks and objects in the environment are unknown to them and can only be inferred from noisy sensory data, leading to considerable uncertainty. Thus, repeatedly walking back to the same target location leads to variability in the endpoints, which depends on the availability and reliability of cues prior to and during the homing response, as well as the subjects’ memory of the target location.

We hypothesized that endpoint variability observed in the homing task reflects the interaction of three sources of sensory-motor noise, namely motor, perceptual, and representation noise (see Figure 2). When executing a goal-directed trajectory in the absence of landmarks, humans rely on path integration (the integration of self-motion cues to determine the position and heading) and an internal representation of the environment [9, 33]. However, path-integration without the correction from external directional references is prone to accumulating noise resulting in increasing positional and heading errors throughout the trajectory [13, 56] (see Figure 2A). Similarly, noise in observation (see Figure 2B) and representation (see Figure 2C) lead to variability in endpoints even in the absence of errors caused by noisy motor actions. As stated above, all three sources of noise interact over time, leading to the observed endpoint variability in homing tasks (see Figure 2D).

**Figure 2.**
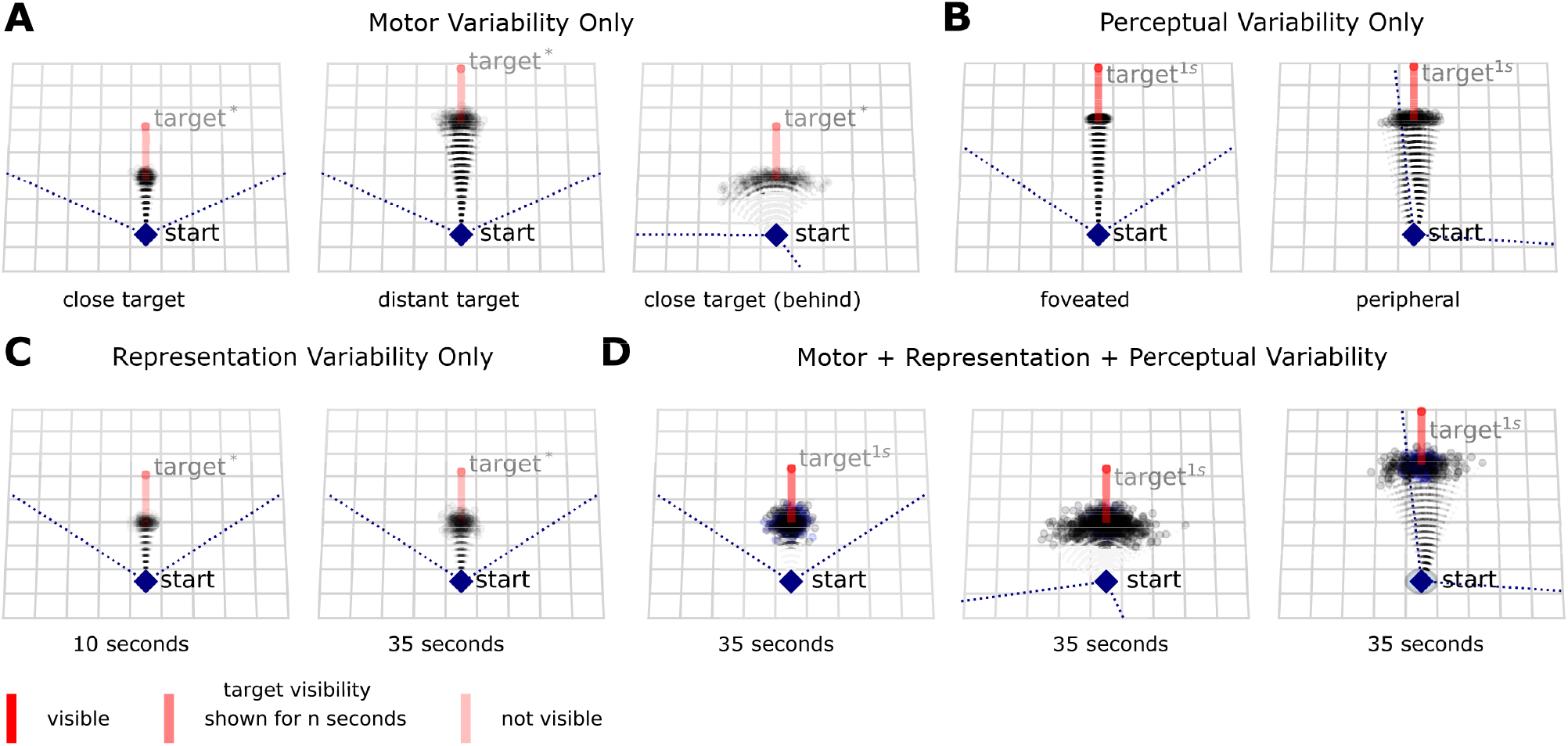
Sources of variability in navigation behavior. (A) Effect of signal-dependent motor noise when walking to a previously seen target. Even when assuming perfect knowledge of the target’s location, different target distances (left, middle) or initial heading directions (right) lead to different patterns of endpoint variability due to signal-dependent noise in linear and angular velocities, which accumulate over the course of the trajectory. (B) State-dependent noisy observations of the same target (visible for 1s) viewed centrally (left) or viewed in the periphery of the visual field (right). Observations are noisier in the periphery compared to the center of the visual field and increasingly so with distance to the observer. When subjects first encounter a landmark, noisy egocentric observations of its distance and direction are used to build up an internal representation [8, 30, 33], which then guides subsequent movement to the (later invisible) target location. (C) Time-dependent noise accrual in internal representation and subsequent movement to the target. Different waiting times after target observation result in different patterns of endpoint variability, even in the absence of noisy motor actions and observations. (D) Combined effect of noise in perception, action, and representation on endpoint variability. Subjects either face the target directly (left), the target is behind them and they store a representation of its location (middle) in memory, or they see the target only in the periphery (right). Blue dots show the contribution of noise in representation to observed endpoint variability.

We propose a computational model of spatial navigation, based on optimal control under uncertainty [51, 52], which considers variability in these three sources as well as behavioral costs for performing motor actions jointly over time. This model allows us to simulate behavior for the entire homing task, including out-bound trajectories (see Figure 1F), as well as subsequent homing under different experimental manipulations. A detailed mathematical description of the different model components, e.g. land-mark perception, internal representation, motor planning and motor action, as well as the choice of model parameters is provided in Methods and Table 2.

**Table 2.**
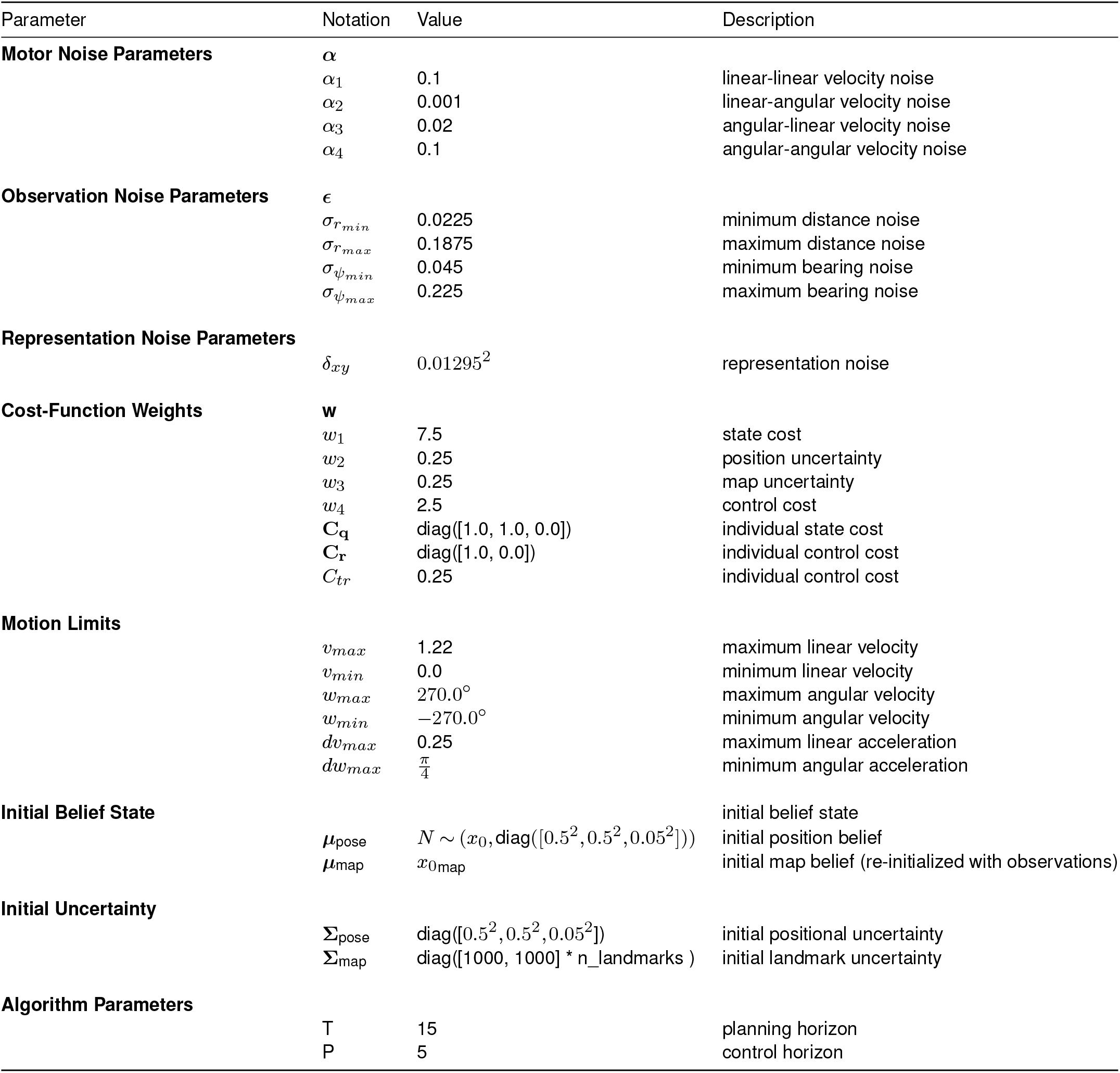
Model parameters for the dynamic Bayesian actor model.

To deal with growing spatial uncertainty, which is the result of noisy motor actions, observations, and internal representations, this model maintains a dynamic belief distribution [47] over possible world states, which serves as the basis for motor planning and is continuously updated by both perception and action [26, 47, 57] (see Figure 3A). Based on this internal belief, motor actions, i.e., forward locomotion and change of heading direction, are selected according to a cost function, which trades off costs for goal reaching, control effort, and uncertainty over the current position as well as the location of landmarks. Throughout the trajectory, noisy motor actions will increase the subject’s positional uncertainty, which is captured by the belief state (see Figure 3B). Over time, uncertainty about the location of goals and landmarks grows in the subject’s representation. When making initial observations of landmarks or objects in the environment stored in the internal representation, subjects consider their current positional and heading uncertainty in relation to the landmark (see Figure 3C). Once a landmark or any object is stored in the internal representation, it provides subjects with orientation and aids in reducing errors caused by sequential noisy motor actions depending on the degree of uncertainty relative to subjects’ positional uncertainty. This allows for beaconing when walking towards visible targets (see Figure 3D) as well as reorientation based on landmarks prior to walking back to the remembered target location (see Figure 3E).

**Figure 3.**
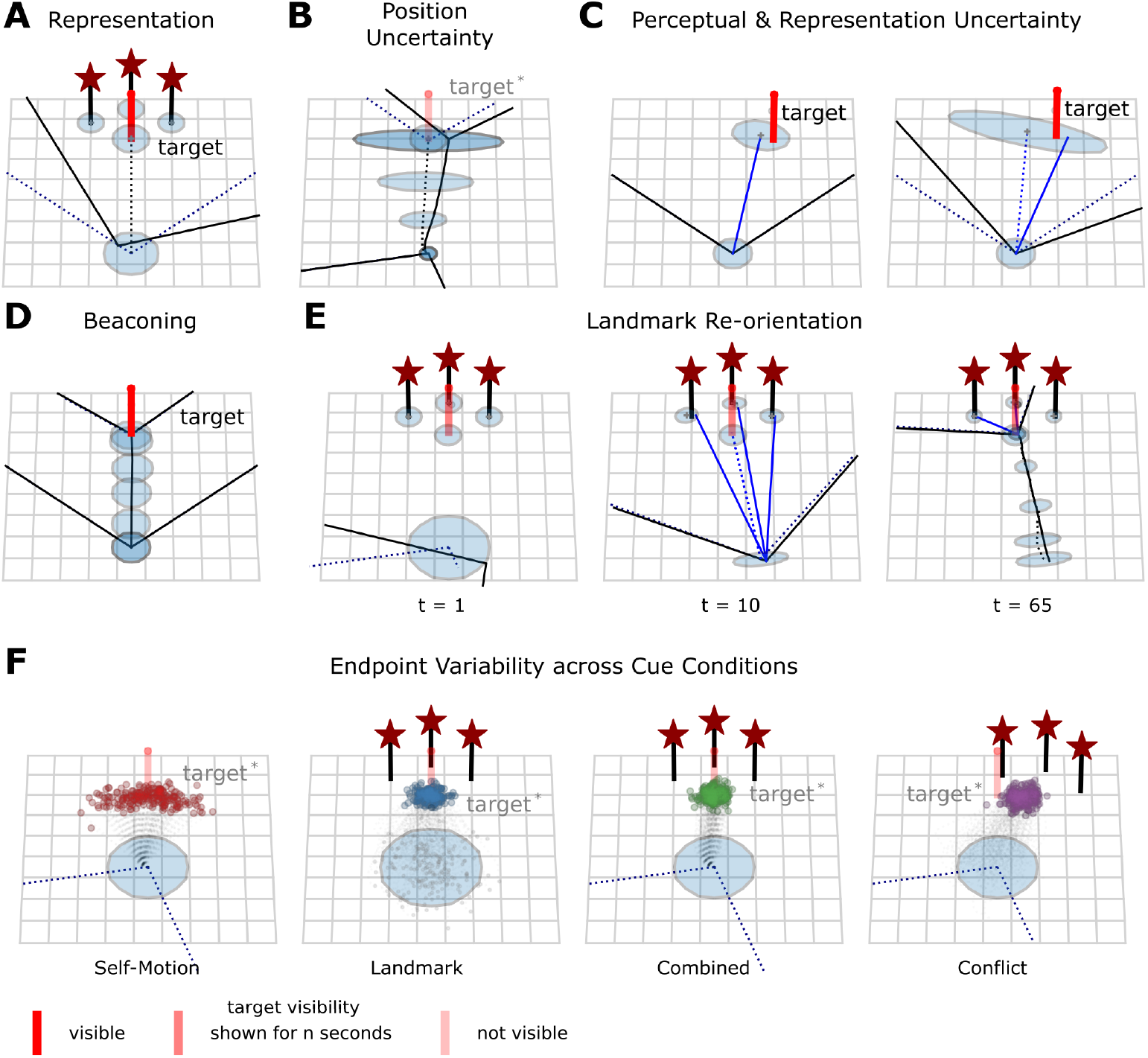
Dynamic Bayesian actor model. (A) Subjects rely on an internal model of their self-location and the location of landmarks in an allocentric reference frame to plan their motor actions. This internal model is updated as a consequence of perception and action. The belief state keeps track of the subject’s spatial uncertainties (blue ellipses & dotted lines). A goal is considered to be reached by the actor when the belief state about self-location matches the belief state of the target location in the internal representation. (B) Noisy motor actions increase positional and heading uncertainty which is tracked by the belief state accordingly. (C) Initial noisy egocentric observations (distance and bearing) are transformed into an internal allocentric representation using the subject’s current positional and observation uncertainties. Left: The subject’s heading direction is fully oriented when observing a landmark. Right: High heading uncertainty in a disoriented subject leads to increased uncertainty about the target’s location when being stored in the internal representation. (D) During goal-directed locomotion towards a visible target, i.e., beaconing, visual feedback from landmarks reduces errors stemming from noisy motor actions. (E) Landmarks aid subjects in resolving positional uncertainty and orienting in the environment. Left: Subject is disoriented, i.e., belief about heading and position (dotted lines) are discrepant from true heading and position (full lines). Middle: Integration of current observations with a prior belief of landmark position resolves positional uncertainty allowing subjects to re-orient. Right: Movement toward the invisible target location using landmark observations to correct errors made by noisy motor actions. Uncertainty about self-location and heading decreases with increasing proximity to landmarks. (F) Dynamic Bayesian Actor model allows simulating homing behavior for different cue conditions [35, 38, 39].

Our dynamic Bayesian actor model allows us to simulate the behavior for different experimental conditions in prior work [35, 38, 39] and obtain trajectories and endpoint variability (see Figure 3F; Methods). We now show that endpoint variability, cueintegration-like behavior, and previously reported seemingly contradictory results [35, 38, 39] under an ideal observer account, are all well captured by a dynamic Bayesian actor model, jointly treating noise in sensory, motor, and representational systems.

### Endpoint variability is explained by noise in perception, action and representation

We simulated 272 trials (16 independent trials for 17 virtual subjects) from our computational model performing the task within the environment from Nardini 2008 [35] under the four conditions (self-motion, landmark, combined, conflict).Figure 4A compares the distribution of endpoints from behavioral data in Nardini 2008 [35] (top) to simulations from our computational model (bottom). Our dynamic Bayesian actor model accounts for the differences across the different cue conditions, as endpoint distributions were not significantly different from the behavioral data (homogeneity energy test between conditions; all p > 0.05; Bonferroni correction applied).

**Figure 4.**
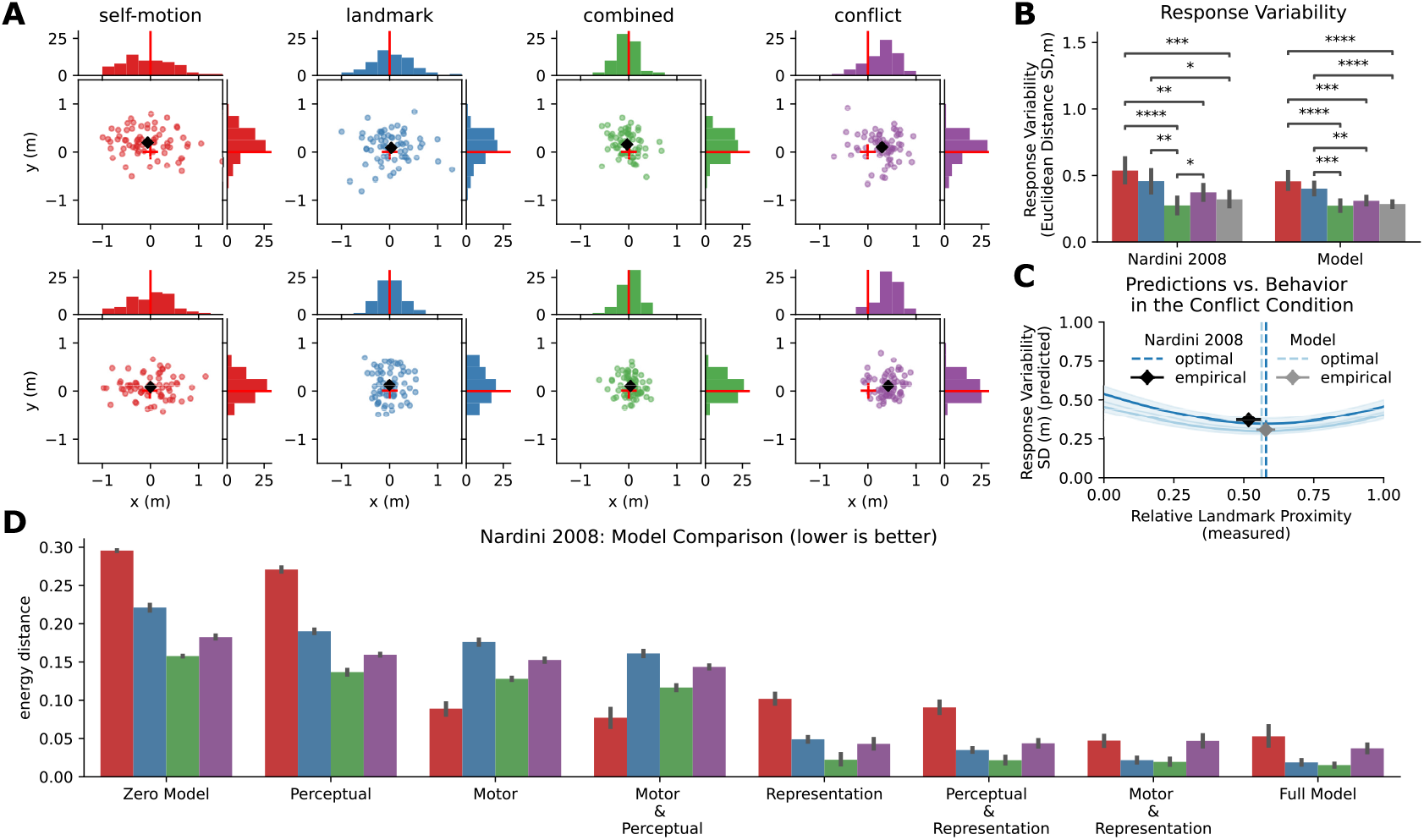
Dynamic Bayesian actor model explaints endpoint variability in a homing task. (A) Endpoint distributions for all four conditions in Nardini 2008 [35]: simulations of dynamic Bayesian actor model (top) and participants’ behavior (bottom). (B) Comparison of response variability of empirical cue-integration account [35] and simulated participants from our Dynamic Bayesian Actor model.*p < 0.05;**p < 0.01;***p < 0.001; ****p < 0.0001; posthoc t-test. (C) Predictions of ideal observer model based on response variability in single cue condition vs. actual behavior in conflict condition. Both mean response variability and mean response proximity across (virtual) subjects are shown for data from Nardini 2008 [35] and simulations from the dynamic Bayesian actor model. Error bars show +-1 standard error of the mean. (D) Comparison of the influence of models’ perceptual, representational, and motor variability on endpoint distribution for the homing task of Nardini 2008 [35]. Energy distances between endpoint distributions of behavioral data vs. model simulations (10 simulations for each model) are computed for each condition. Errors bars indicate the standard deviation of energy distances. Full endpoint distributions are shown in Figure S2.

We were further interested in the response variability for each of the four conditions and predictions of the optimal cue integration model [35].Figure 4B shows observed standard deviation in homing response (response variability) similar in magnitude for data by Nardini 2008 [35] and simulations from our dynamic Bayesian actor model. A two-way repeated measures ANOVA on response variability with type (model simulation vs. empirical data) and cue condition (self-motion, landmark, combined, conflict) as independent variables revealed a significant difference for condition (F(4,120) = 25.83, p < 0.05), but not for type (F(1,32) = 2.88, p = 0.1) as well as no significant interaction effect for type and condition (F(4,128) = 0.7, p = 0.6). Using the ideal observer model, we predicted homing variability in the double cue condition, i.e., combined and conflict, using the observed variability from single cue conditions (see Figure 4C; green vs. gray bars). Both human and model behavior agree with predictions made by the ideal observer model, which predicts that variability in the combined cue condition is lower than in both single cue conditions (all p < 0.05). Our dynamic Bayesian actor model also predicts subjects’ behavior in cue conflict conditions in terms of response proximity to landmark cues following predictions made by the ideal observer model (see Figure 4C).

Based on our hypothesis that endpoint variability reflects the interaction of three different sources of sensory-motor noise, we tested the contribution of perceptual, representation, and motor variability separately by formulating eight different models (see Methods). We compared endpoint distributions generated by these models against behavioral data (see Figure S2 for simulated endpoint distributions).Figure 4D shows energy distances between the empirical distribution and simulated distributions for all models and all conditions. These results show that neither motor, perceptual, nor representation variability alone are sufficient to account for the observed endpoint variability of subjects across all four experimental conditions. Subjects’ memory for landmark and target locations had the strongest influence on overall end-point variability across cue conditions, whereas the influence of perceptual variability was negligible. This is most likely, because subjects observed the entire environmental layout, including the locations of all three targets and the three landmarks in the background at the beginning of each trial, together with the three landmarks in the background,. This possibly allowed subjects to form a comparatively accurate representation of the environment. Further, the contribution of motor variability was highest in the selfmotion condition, in which no landmark information was available during homing.

This result establishes that a dynamic Bayesian actor model incorporating sensory, representational, and motor uncertainty is able to capture navigational variability observe in the homing task with landmarks across different cue conditions during homing.

### Optimal Control under perceptual, motor & representational uncertainty predicts behavior across studies and experimental conditions

We then asked whether the dynamic Bayesian actor model can predict endpoint distributions and endpoint variability across different homing tasks with different environmental layouts, i.e., differing in locations of goals as well as the number and locations of landmarks. To test this, we simulated trials using the environments (see Figure S1) and task parameters (see Table 1) of Chen 2017 [39] (Experiment 1a: one vs. three landmarks) and Zhao 2015 [38] (proximal vs. distal landmarks). We kept all model parameters of the dynamic Bayesian actor constant to assess how well our model predicts observed variability across different homing tasks with differing experimental manipulations.

To obtain endpoint distributions in the different cue conditions, we simulated 1440 trials (18 independent virtual subjects with 40 trials for each experiment) from experiment 1a in Chen 2017 [39], in which humans performed the homing task in either an environment with three landmarks (rich) or only a single landmark (poor). Differences in terms of the distribution of endpoints between the two environments (one vs. three landmarks) and the four cue conditions were well predicted by the dynamic Bayesian actor model (see Figure 5A and B). Additionally, we computed response variability within all four conditions for both environments and compared simulations of the dynamic Bayesian actor model against empirical data (see Figure 5C and D). Response variability across cue conditions was again similar between behavioral data and simulated data. Reduction of response variability in double cue conditions, compared to single cue conditions, matched optimal predictions of the ideal observer model in both environments (see Figure 5C and D; green vs. gray bar).

**Figure 5.**
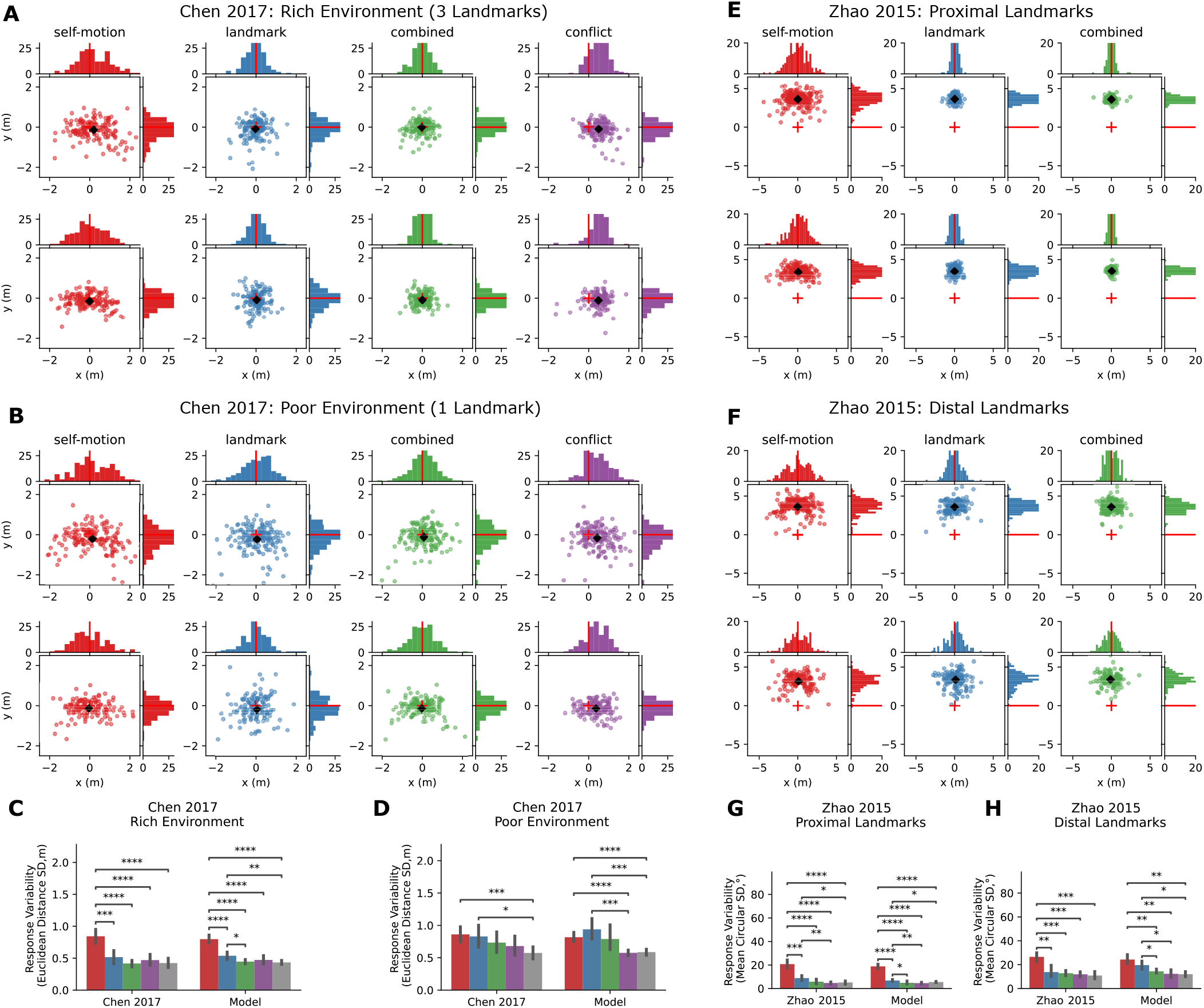
Dynamic Bayesian actor model explains endpoint variability across studies & environments. (A and B) Endpoint distributions from Chen 2017 [39] (top; A: Exp1a rich environment; B: Exp1a poor environment) and model simulations from dynamic Bayesian actor model (bottom; A: Exp1a rich environment; B: Exp1a poor environment). Endpoint distributions are not significantly different between our model and the empirical data for the environment with three landmarks (homogeneity energy tests between conditions; all p > 0.05; Bonferroni correction applied) and the environment with one landmark (homogeneity energy test between conditions; all p > 0.05; Bonferroni correction applied). (C and D) Response variability (euclidean distance) across cue conditions for Chen 2017 [39] (C: Exp1a rich environment; D: Exp1a poor environment) and model simulations from dynamic Bayesian actor model. In the environment with three landmarks, we found a significant main effect of cue condition (F(4,136) = 75.22, p < 0.001), no significant main effect of type (F(1,34) = 0.01, p = 0.93), and no significant interaction effect between cue condition and type (F = (4,136) = 0.59, p = 0.67). Similarly, in the environment with a single landmark, we found a significant main effect of cue condition (F(4,136) = 12.36, p < 0.001), no significant main effect of type (F(1,34) = 0.01, p = 0.94) and no significant interaction effect between cue condition and type (F = (4,136) = 1.28, p = 0.28). *p < 0.05;**p < 0.01;***p < 0.001; ****p < 0.0001; post-hoc t-test. (E and F) Endpoint Distribution from Zhao 2015 [38] (top; E: proximal landmarks; F: distal landmarks) and model simulations from dynamic Bayesian actor model (bottom; E: proximal landmarks; F: distal landmarks). Data for conflict condition are shown in Figure S5 (G and H) Response variability (homing direction) across cue conditions for Zhao 2015 [38] (G: proximal landmarks; H: distal landmarks) and model simulations from dynamic Bayesian actor model. In the environment with proximal landmarks, we found a significant main effect of cue condition (F(4,40) = 81.91, p < 0.001), no significant main effect of type (F(1,10) = 3.23, p = 0.1), and no significant interaction effect between cue condition and type (F(4,40) = 0.59, p = 0.67). Similarly, in the environment with distal landmarks, we found a significant main effect of cue condition (F(4,32) = 31.37, p < 0.001), no significant main effect of type (F(1,8) = 0.59, p = 0.46) and no significant interaction effect between cue condition and type (F = (4,32) = 2.05, p = 0.11), suggesting that response variability was similar between model simulations and empirical data across the four experimental conditions and the two landmark environments. *p <0.05;**p < 0.01;***p < 0.001; ****p < 0.0001; post-hoc t-test.

We also simulated data from two experiments from Zhao 2015 [38], in which landmarks were either 5.5m away (proximal landmarks; 2160 trials for 6 virtual subjects; 360 trials per subject) or 500m away in the same relative configuration (distal landmarks; distal landmark group - 1800 trials for 5 virtual subjects; 360 trials per subject).Figure 5E and F show endpoint distributions for both the proximal and distal landmark environments. Visual inspection yields similar endpoint distributions between empirical data and model simulations for proximal and distal environments. Although qualitatively in agreement, quantitative comparison using homogeneity energy tests for differences between both empirical and model endpoint distributions exhibited differences in both proximal and distal environments (all p < 0.05; Bonferroni correction applied). Statistically significant differences between the endpoint distributions are most likely due to a larger sample size (3 and up to 7 times as much data than in other experiments). Based on the subjects’ endpoint distributions, we computed subjects’ heading errors relative to the correct mean response direction for each of the four target locations. Analysis of response variability shows heading errors, which are similar in magnitude between empirical data and model simulations for both the environment with proximal landmarks and the environment with distal landmarks (all p > 0.05, see Figure 5G-H).

Unlike in the experiment performed by Nardini 2008 [35] in which the environment was fully observable at the beginning of each trial, target locations in these four experiments appeared sequentially, requiring subjects to learn the environmental layout during the unfolding of the triangle completion task. When comparing models differing in the availability of the perceptual, representational, and motor noise, perceptual and motor noise had a relatively higher contribution to overall endpoint variability across the different homing conditions (see Figure S3 and Figure S4) compared to the environment from Nardini 2018. However, representation variability remained the largest influence across the four different conditions. and models including both representation and motor variability provided the closest matching endpoint distributions.

Taken together, our computational model is able to explain data across a multitude of experimental conditions [35, 38, 39] with varying configurations of landmarks, environmental layouts, and walking distances under the same set of model parameters. We now show that our dynamic Bayesian actor model can explain behavior seemingly sub-optimal according to a cue-integration model as optimal sequential actions when considering perceptual, motor, and representational uncertainties jointly.

### Optimal sequential actions under uncertainty predict seemingly sub-optimal ‘cue weights’ in conflict conditions

One previous result by Chen 2017 [39], which the cue integration model cannot explain, concerns the discrepancy between optimal ‘cue weights’ (computed from the variances of walking endpoint distributions in single cue conditions) and empirical ‘cue weights’ (computed from the means of response distributions). Based on the observed response variability in single-cue conditions, Chen 2017 computed optimal ‘visual cue weights’, which, according to a perceptual cue integration account, should be reflected in subjects’ reliance on landmark cues compared to selfmotion cues in the two double cue conditions, i.e., combined and conflict. Predictions of such optimal ‘visual cue weights’ are explicitly tested in conflict conditions by rotating landmarks 15°around the location of the last target. This allows for determining whether subjects’ responses follow the self-motion or the landmark-consistent location, or fall somewhere between the two. Based on response proximity to either cue (see Figure 6A left), these empirical ‘visual weights’ are computed as the relative response proximity to either cue.Figure 6B (left) shows the linear relationship between both types of ‘cue weights’ for subjects in the two environments for data of Chen 2017 [39]. Analysis of regression coefficients shows intercepts larger than zero and slopes less than one, indicating a regression to the mean (poor environment: R^2^ = 0.361, p = 0.011, slope 95% ci (0.099, 0.636); rich environment: R^2^ = 0.45, p = 0.003, slope 95% ci (0.195, 0.804)). According to the cue integration interpretation [39, 44], subjects underweighted the influence of self-motion cues and overweighted the influence of landmark cues, as measured by the response variability in both single cue conditions.

**Figure 6.**
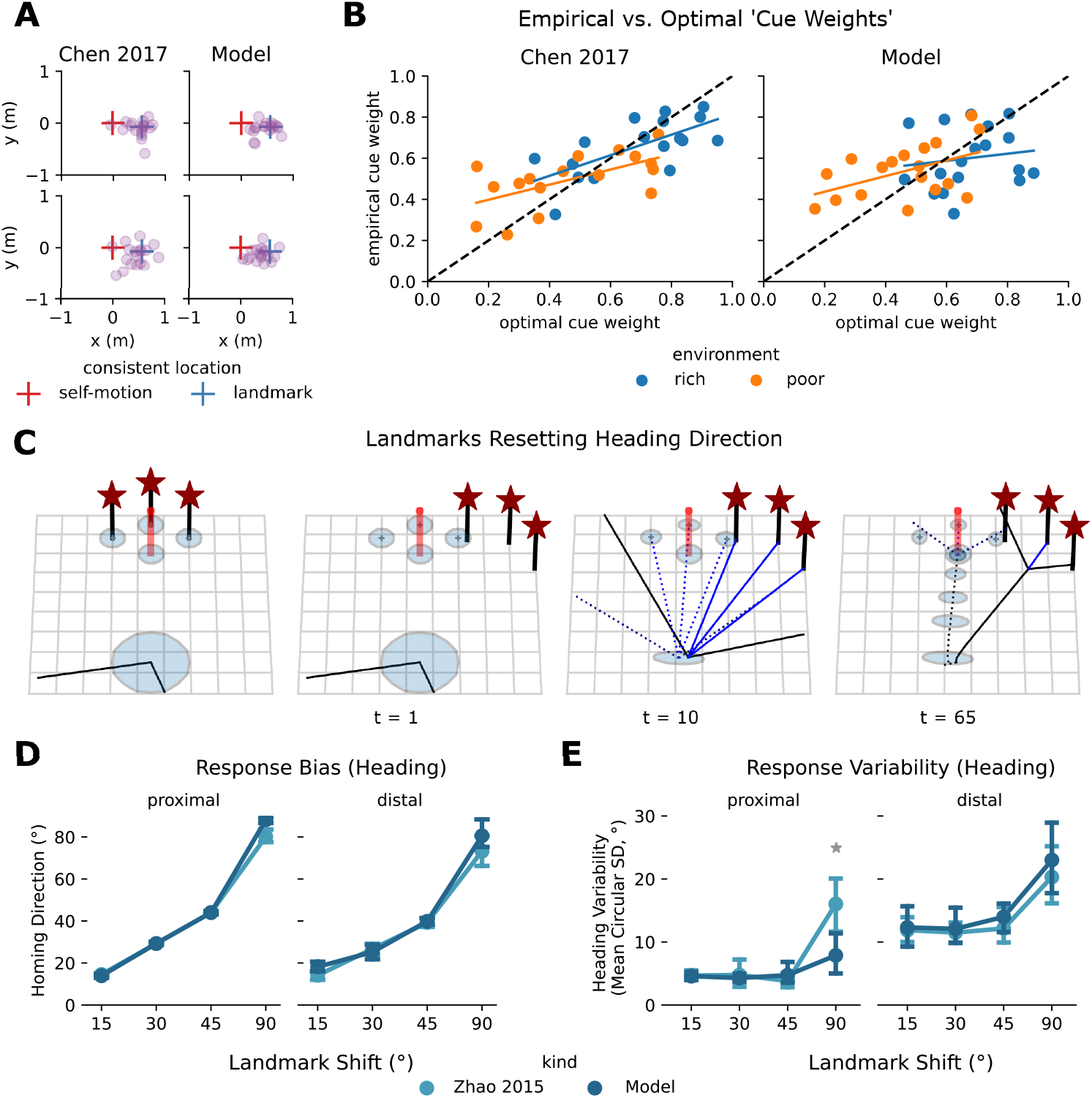
Optimal sequential actions under uncertainty predict seemingly sub-optimal cue-integration in conflict conditions. (A) Mean responses (single cue condition bias corrected) for conflict condition for data from Chen 2017 Exp1 [39] (left) vs. dynamic Bayesian actor model (right) for rich (top) and poor environments (bottom). (B) Optimal vs. empirical cue weights for data of Chen 2017 [39] vs. our dynamic Bayesian actor model. Dots indicate individual subjects. Optimal cue weights were computed based on observed response variability, whereas empirical cue weights were computed based on response proximity in conflict conditions to either cue’s location. Linear Regression lines were fitted to subjects’ cue weights for the two environments, poor and rich. (C) Dynamic Bayesian actor explains the dominance of landmarks cues in homing direction. 1: Subjects maintain a belief about self-location, as well as the location of landmarks. 2: Landmarks are covertly rotated, whereas the belief about landmark locations, i.e., their expected position, remains unchanged. 3: The subject turns around, bringing the landmarks into the field of view while also increasing uncertainty about the heading direction due to the rotation. Egocentric observations of the landmarks (distance and bearing) reset the heading of the subject because the subject expects the landmark to the direction straight ahead. Uncertainty in position and heading is reduced. 4: For homing, subjects utilize the underlying belief-space representation to navigate towards the final goal, using the landmarks to correct any motor errors, further reducing positional uncertainty, this leads to lower endpoint variability. (D) Response bias for conflict conditions from Zhao 2015 [38] vs. model behavior from dynamic Bayesian actor model. Endpoint distributions are shown in Figure S5 (paired t-tests; all p > 0.05; Bonferroni correction applied). (E) Response variability for conflict conditions from Zhao 2015 [38] vs. model behavior from dynamic Bayesian actor model (paired t-tests; p > 0.05 for angles of 15°, 30° and 45°; Bonferroni correction applied).

Here, we hypothesized that such seemingly sub-optimal ‘cue weights’ arise as optimal actions under perceptual, motor, and representational uncertainties from the perspective of our dynamic Bayesian actor model, rather than making additional assumptions about cost-functions, penalizing responses deviating from the mid-point between the two cues in to correct these seemingly sub-optimal ‘cue weights’ [44]. Based on response variability (shown in Figure 5C-D) and mean responses in conflict conditions (shown in Figure 6A; right) we computed both types of ‘cue weights’ in the two environments, replicating the analysis from Chen 2017 [39] for virtual subjects from our computational model.Figure 6B shows empirical versus optimal ‘cue weights’ in subjects from Chen 2017 [39] (left) and virtual subjects from our dynamic Bayesian actor model (right) for the poor and rich environment. Visual inspection yields that optimal vs. empirical ‘cue weights’ are similar in magnitude between our computational model and empirical data from Chen 2017. One exception is that data from Chen 2017 exhibited larger variability between subjects’ optimal ‘cue weights’, which was not entirely captured by our computational models’ virtual subjects, since they were all simulated from the same underlying model parameters. Linear regression analysis showed slopes smaller than one and intercepts larger than zero (poor environment: R^2^ = 0.230, p = 0.044, slope 95% ci (0.011, 0.751); rich environment: R^2^ = 0.021, p = 0.576, slope 95% ci (−0.452, 0.796)), indicating a regression to the mean effect, as reported in Chen 2017 [39].

Taken together, seemingly sub-optimal cue integration behavior is well captured by the dynamic Bayesian actor model considering variability in perception, action, and representation jointly. Notably, the present analysis demonstrates that endpoints of walking trajectories are better accounted for as dynamically arising from the interaction of perception, internal representations, and action instead of as static perceptual ‘cue weights’.

### Response variability is reduced, while a single cue predicts response direction

For conflicts larger than 15°, Zhao 2015 [38] observed an incongruity between heading variability and heading direction that could not be attributed to a previously proposed ideal observer model [35]. It was found that response direction was almost entirely dominated by a single cue, i.e., the shifted landmarks, while response variability was reduced simultaneously, violating the optimal cue integration prediction [43]. If subjects had integrated these cues optimally according to perceptual cue integration, their response direction would have been a weighted average according to the relative reliabilities observed in single cue conditions. Instead, subjects seemed to rely on a single cue, while response variability was reduced when navigating towards the shifted target’s location with landmarks in sight during homing.

Our dynamic Bayesian actor model offers an alternative view of this behavior by jointly considering the effects of perception, action, and representation during homing behavior.Figure 6C illustrates how landmarks orient subjects by resetting their heading direction in the internal representation for a conflict of 30°. During subsequent homing, the landmarks aid subjects in reducing positional and heading uncertainty, which would otherwise accumulate, leading to reduced response variability, although homing responses are biased. The simulations demonstrate this behavior for conflict conditions of varying degrees between 15°and 90°found in Zhao 2015 [38] (proximal and distal landmark environments; see Figure S5). Figure 6D shows how the dominance of shifted landmarks in subjects’ response direction is well captured by our dynamic Bayesian actor model for both the proximal (left) and distal landmark (right) environments. Moreover, the reduction in response variability was also predicted by our dynamic Bayesian actor model across the different landmark shift conditions (see Figure 6E)

We see a discrepancy at a conflict angle of 90° between our model and behavioral data (paired t-test; p = 0.01; Bonferroni correction applied), as subjects in Zhao 2015 [38] seemed to notice the conflict between response directions on some trials. Thus, they ignored landmarks cues altogether in those trials leading to increased response variability. This was generally the case for conflicts larger than 90°, hence they were excluded from the present analysis. Such switches in behavior can, in principle, be modeled by additional assumptions about trial-by-trial learning and decision rules for the large discrepancy between the expected position of landmarks given subjects’ uncertain internal belief about their heading direction and their noisy observed heading direction, e.g., causal inference [58].

## 3 Discussion

Navigating from one place to another requires utilizing information from spatial cues, both internal to the actor and external from the environment, representing them internally, planning and executing motor actions sequentially. A popular family of tasks that have been used to investigate the contributions of different sources of information to spatial navigation behavior are triangle-completion (or homing) tasks [3]. While prior research has shown that humans are able to combine different spatial cues and thereby reduce their navigational variability [34, 35, 37, 39–41, 44, 59, 60], it is still unclear how or why the observed variability in trajectories and endpoints arises and how path-integration and landmark cues interact in this process, which necessarily requires recurring to a computational model.

Here, we hypothesized that the spatial uncertainty about task-relevant quantities in the internal model of the navigator, either about self-location, heading direction, or the location of landmarks and objects within the environment, interacts dynamically with different cues during navigation and results in the observed endpoint variability in homing tasks. Because a vast array of natural sensorimotor behaviors in humans, including basic motor control tasks [46, 61–63], locomotion [64] and ball-catching behavior [48], has been explained successfully in terms of optimal control under uncertainty [51], we formulated a dynamic Bayesian actor model of human navigation. This model provides a unifying account of endpoint variability and explains several seemingly contradictory phenomena observed in previous work. Simulations of the homing tasks from different studies show endpoint distributions in different cue conditions closely resembling those of human participants. Previous ideal observer models of navigation [35, 38, 39], include no actions and therefore cannot explain how endpoint variability arises in single cue conditions, instead relying on empirically measured variability to predict optimal behavior in double cue conditions.

Empirical data were analyzed with model behavior obtained from simulations by the dynamic actor model and under an ideal observer account for spatial navigation [35]. Quantitative analysis of endpoint distributions by computing response variability, either over euclidean distance [35, 39] or heading direction [38] to mean response locations, shows response variability and proximity similar in magnitude between empirical data and model simulations across different homing tasks and experimental manipulations. Selectively excluding individual sources of uncertainty provided consistently larger deviations in the endpoint distributions, demonstrating the necessity to include all uncertainties and their interaction. The dynamic Bayesian actor also predicted behavior in conflict conditions, considered sub-optimal under the ideal observer account. Thus, we find strong evidence that human navigation behavior in triangle completion tasks needs to take the different contributions of uncertainty from perceptual, motor, and representation systems into account, which can be well captured by optimal control under uncertainty.

### Relation to other experimental work

Beyond being able to capture and explain empirical data from multiple triangle completion tasks, including behavior considered contradictory under ideal observer models, the dynamic Bayesian actor model allows to reconcile previous accounts of navigation in a coherent computational framework. The present model includes perception [8], representation [33], planning [65] and execution of motor actions [12, 13, 56], while considering uncertainty, which allows predicting and explaining how these seemingly separate processes interact in producing observed navigation behavior in triangle completion task.

The homing task with landmarks has been described as comprising several different ‘navigational strategies’ while the task unfolds. Our dynamic Bayesian actor model dynamically shifts between these strategies implicitly, depending on the availability of different spatial cues and their relative uncertainty. Strategies that the dynamic Bayesian actor model shows include walking toward a visible target location, i.e., beaconing [29], and continuously updating self-location based on internally generated motor cues, i.e., path-integration [11]. The presence of landmarks reduces motor errors and, thus, endpoint variability by orienting subjects in the environment [25] and continually providing visual feedback about their location. Here, cue integration happens implicitly and sequentially but accounts only for state estimation. In this view, landmark perception effectively ‘corrects’ whatever errors arise when executing noisy locomotor actions, thus relating path-integration to landmark-based navigation strategies [26].

The present model can also explain previous data such as humans walking in circles in the absence of landmarks [13] and reorienting in the presence of known landmarks [25]. Furthermore, the underlying internal map explicitly represents landmark uncertainty, which mediates how much subjects should rely on landmark cues for determining their position and heading direction. For example, in environments with low environmental stability, humans have been shown to rely less on landmarks and more on path-integration cues [39, 59, 66], a phenomenon readily explainable when considering the interaction of uncertainties. Finally, the dynamic Bayesian actor further offers a computational level explanation of the role that path integration plays in building up an allocentric cognitive map from sequential egocentric observations of landmarks and objects [10, 19, 28, 67] and how this internal map is used subsequently to plan and execute movements [33].

### Relation to other computational models

A vast array of previous computational models of sequential goal-directed navigation behavior have been based on differing approaches, including artificial neural networks [68–70], reinforcement learning [65, 71–73], or a combination thereof [23]. Furthermore, individual modeling solutions can be distinguished by the respective navigation tasks considered, their complexity, the actions, and state information available to agents, which fundamentally influences the learning of internal representations, self-localization, and planning. Importantly, in complex real-world navigation, the position and heading of the agent in relation to the environment can never be sensed directly, making it both ambiguous and uncertain. This implies that the current state from the agent’s perspective can only be inferred based on sequences of observations together with a latent representation of the environment which has to be learned upon encountering a novel environment [26, 67]. Thus, an important feature for distinguishing computational models is their ability to correctly handle the different sources of uncertainty.

Investigations involving model-free and model-based reinforcement learning in virtual mazes have yielded that human behavior is consistent with a model-based strategy [65], i.e., planning. More recent work explored the possibility that humans may use a hybrid strategy between model-free and model-based approaches [72–74]. However, in these maze environments, it is often assumed that there is no perceptual uncertainty about the state, i.e., the current state is fully observable, the actions available to the agent are a small set of discrete choices, and variability is associated only with learning across episodes or switching between model-based and model-free strategies. In principle, maze-like environments can be altered by introducing locations that look the same, requiring the agent to perform localization based on the history of previously visited locations [75, 76].

Neural networks models of spatial navigation range from models specifically trained to perform a particular task in an end-to-end fashion [23, 70, 77] to more descriptive and mechanistic models taking into account known biological constraints about the underlying neural circuits in the hippocampal formation [68, 69]. Artificial neural networks trained on the task of localization in openfield environments, i.e., determining position from inputs of noisy linear and angular velocities, have exhibited grid cell-like activity hidden units [23, 77] suggesting that the brain might solve a similar optimization problem. However, the emergence of grid cells is highly sensitive to particular hyper-parameters and implementation details [78]. The emergent grid-like representation can be combined with a network providing intermittent visual input from external landmarks to perform more accurate localization and a policy network to facilitate navigational planning [23]. The resulting system successfully navigates towards remembered goal locations while taking shortcuts, in both open-field and maze environments, suggesting an integral role of grid cells for vector-based navigation. However, none of these neural network models can handle the sequential integration of uncertainties stemming from perception, internal models, or actions.

Modeling the triangle completion task with landmarks requires solving the computational problems of spatial localization, learning of an internal representation, planning, and control in a partially observable environment. Bayesian methods allow to capture and reason about this uncertainty in a principled manner allowing for a combination of different cues or the use of prior knowledge to reduce uncertainty and, thereby, variability. In order to solve this problem sequentially, our dynamic Bayesian actor model employs a dynamic belief-space representation [26, 47] in a continuous control task with internal model uncertainty. This dynamic belief representation is general enough to be integrated with other probabilistic Bayesian accounts of spatial navigation. Besides the interaction of landmark and path integration studied in the present work, Bayesian cue integration processes have been shown to be important for integrating linear and angular velocities from different sources, such as visual, vestibular, and proprioceptive cues, in order to reduce positional uncertainty and bias [79]. Furthermore, the probabilistic nature of our model lends itself to incorporating biases in self-motion perception, e.g., slow-velocity priors [80], or priors over target locations [81] as well as studying spatial learning across trials [81, 82], the monitoring of environmental stability [39, 59, 66] and causal inference [34, 58] to explain behavior for highly discrepant cues.

### Relation to neuroscience

The dynamic Bayesian actor model presented here provides computational-level explanations for the observed behavior, allowing to relate behavior to internal representations and neuronal computations [83, 84]. Computationally speaking, the neural substrate underlying spatial navigation, in particular, places cells in the hippocampus [18], grid cells in the medial entorhinal cortex [85] and head-direction cells in the postsubiculum [86], serve as an internal model of the surrounding environment in relation to the actor. This internal model is sequentially updated by both internally generated self-motion cues [19, 87, 88] and external cues [89–92], supports learning and adaption in novel environments [67, 93, 94] and is involved in the planning of future actions [23, 65, 95, 96]. Thus, the present dynamic Bayesian actor model provides a normative computational model of the subjective internal beliefs and their sequential evolution, which can be related to neuronal activities.

Path integration is supported by an interaction of place, grid, and head direction cells, which enables self-localization relative to an internal allocentric representation [19, 88], even in the absence of external cues [85, 97]. Beyond self-localization based on internal cues, the dynamic Bayesian actor implies at least two other key roles of path integration in relation to an internal cognitive map. Namely, path-integration is involved in linking together sequences of egocentric observations into a unified allocentric representation, i.e., cognitive mapping [67], and in determining self-location and heading based on sequences of observation and prior predictions based on the internal representation, i.e., reorientation [25, 26]. The internal map is utilized for goal-directed planning of movement [23, 65, 67, 95], which is then transformed into motor output [96].

However, due to noisy self-motion signals during path integration, the internal representations found in place, grid, and head direction cells may not necessarily correspond to the actual physical location and orientation in the environment [98]. The internal representation rather expresses the current belief regarding these quantities [47], which may become increasingly discrepant from the actual state of the world over time, leading to ever-increasing spatial uncertainty and consequently navigational variability [27]. In order to correct for this discrepancy, the brain needs an error correction mechanism based on stable and more reliable cues, e.g., landmarks or boundary information [90–92] to recover position and heading relative to the internal representation [25, 28].

As perceptual, representational, and motor processes are inevitably noisy, our model may facilitate understanding of how the resulting uncertainty influences internal representations and, consequently, observable behavior. Theoretical work suggests that place and grid cell representations may have evolved to be tolerant to sensory-motor noise [27, 99–101], however it remains unclear to what degree the resulting uncertainty is explicitly represented in the nervous system [102, 103]. Several observations and theoretical accounts suggest a potential role of uncertainty in spatial representations [26, 104–109]. For example, the degree to which landmarks facilitate reorientation depends on their perceived stability [110, 111], implying that the brain learns and keeps track of environmental stability over time. Recent work suggests that the spatial uncertainty may also be encoded in the distorted or enlarged firing patterns of place and grid cells in novel environments [104, 105, 107]. For example, mapping of external space occurs at different spatial resolutions depending on the availability of local visual cues such as landmarks [107]. In particular, place fields coding for particular locations are smaller in environments that provide local visual cues, i.e., they have higher resolution, compared to place fields in larger and more sparse environments, which exhibit larger place field size [107]. Similarly, larger place fields have been observed as a function of distance to boundaries in the environment [106]. As our model explicitly accounts for the uncertainties and variability involved in navigation, it provides a computational basis for relating subjective internal beliefs to neuronal computations.

## Conclusion

Taken together, optimal control under uncertainty provides a unifying computational account of the different sources of noise and their interaction in navigation and explains the observed behavior of human subjects across multiple homing tasks. The dynamic actor considers the interconnected nature of navigational problems, including state estimation, control, learning, and planning, while also accounting for uncertainty in perception, action, and representation over space and time. Our dynamic Bayesian actor model not only explains behavior previously deemed contradictory under ideal observer models but will provide invaluable insights for future work in neuroscience aimed at relating neuronal activity to internal perceptions instead of external stimuli [83, 84], and in the behavioral sciences, as it makes testable predictions for behavior in spatial navigation tasks [112]. The proposed model allows generating new hypotheses and experiments to provide further insights into the computational processes underlying human spatial navigation, a natural task of fundamental importance.

## Acknowledgements

We thank Marko Nardini, Xiaoli Chen, Timothy McNamara, Mintao Zhao and William H. Warren for providing their behavioral data and further clarification regarding the experimental and analysis procedures. Calculations for this research were conducted on the Lichtenberg high performance computer of the TU Darmstadt. This research was supported by the European Research Council (ERC; Consolidator Award ‘ACTOR’-project number ERC-CoG-101045783). We additionally acknowledge support by ‘The Adaptive Mind,’ funded by the Excellence Program of the Hessian Ministry of Higher Education, Science, Research and Art.

## Methods

### EXPERIMENTAL MODEL AND SUBJECT DETAILS

#### Homing Task Description

On each trial, subjects viewed the environment from their starting position. They were then asked to walk an outbound path through a sequence of three goal locations defining a three-legged outbound path. Goal locations were presented visually simultaneously from the beginning of the trial [35] or in succession after completion of each paths leg [38, 39]. Once subjects reached the location of the last goal, the environment turned dark. After a brief delay of 8-20 seconds, the environment turned visible again. Internal or external cues were manipulated under one of four cue conditions. In self-motion conditions, landmarks were removed from the environment. In landmark conditions, subjects were disoriented by placing them in a swivel or wheelchair, moving their position, and turning them at a particular speed for the waiting time. In combined conditions, subjects stood still for the duration of the waiting time, and landmark cues were available during the homing response. In conflict conditions, landmarks were rotated clockwise or counterclockwise around the location of the last post, creating a conflict between self-motion and landmark-defined home locations. Subjects then had to walk back to the (remembered) location of the first target, i.e., homing, utilizing both internal and external cues depending on the cue condition. Response locations were given by their stopping location, i.e., where subjects believed they had successfully reached the home location, for each trial. Depending on the cue condition (self-motion, landmark, combined, conflict), different patterns of endpoint variability are observed.

#### Experiment & Dataset Description

We obtained endpoint data from three studies investigating the homing task with landmarks. Within each of the datasets, we selected data from specific experiments differing in their environmental complexity. These experiments were namely adults performing the homing task (Nardini 2008 [35]), comparison of number landmarks in the environment (Exp1a: poor vs. rich environment; Chen 2017 [39]) and adults performing the homing task in an environment with either proximal or distal landmarks (Zhao 2015 [38]). The environments between the three studies differed in size, the location of targets and landmarks, and subjects’ starting positions. Landmark locations remained stable across trials, while the configuration of target location was varied across trials, leading to differently shaped three-legged outbound paths (see Figure S1A). In virtual environments, i.e., the studies of Zhao 2015 and Chen 2017, subjects wore a head-mounted display that considerably restricted their field of view. The tasks between the studies further differed in the disorientation procedure within the landmark condition, i.e., speed of turning, final rotation, and positional offset), as well as the waiting time at the end of the outbound path prior to the homing response. We extracted the differing task, subject, and environmental parameters from the respective papers and the accompanying trajectory data (when available) (see Table 1).

#### Ideal Observer Analysis

We adopt the notation by McNamara 2021 [44] to describe ideal observer analysis for the homing task. Subjects infer a target’s location *L*, given self-motion *S* and visual landmark cues *V*. Using Bayes rule, the posterior distribution over target locations given visual and self-motion can be written as:

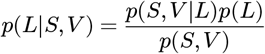

where *P*(*L*) refers to the prior distribution over possible target locations. Assuming conditional independence between self-motion and visual landmark cues, this is expressed as follows:

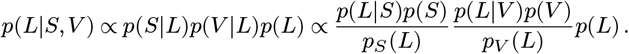

Assuming non-informative priors for location L and both types of cues S and V, i.e., *p*(*L*) = *p*(*S*) = *p*(*V*) = 1, the expression further simplifies to:

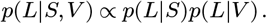

In the cue integration account, subjects weigh self-motion cues *p*(*L*|*S*) and landmark cues *p*(*L*|*V*) according to their relative reliability. Cue reliability is obtained from response variability in single cue conditions by eliminating the influence of the other cue experimentally prior to the homing response either removing landmarks (self-motion condition) or disorienting subjects via turning (landmark condition).

#### Ideal Observer Analysis (Euclidean Distance)

In order to predict subjects’ response variability and bias in the double cue condition, we computed response variability in each cue condition as the euclidean distance for each response to the mean response location. For the analysis, all error distributions in single cue, i.e., *p*(*L*|*S*) and *p*(*L*|*V*), and double conditions, i.e., *p*(*L*|*S,V*), are assumed to be normally distributed (self-motion: 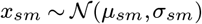; landmark: 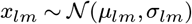; combined: 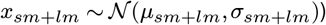.

The assumption of gaussian error distributions allows for a weighted linear combination of the two single cue gaussian distributions relative to their response variability to predict response variability and bias in double cue conditions [35, 39]. Based on the observed response variability in single cue conditions, optimal ‘cue weights’ (*w_lm_* + *w_sm_* = 1) can be computed as follows:

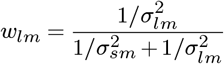

The variance in double cue conditions can be predicted from response variance in single cue conditions as follows:

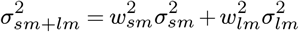

This weighted average yields lower response variability in double cue conditions compared to the variance of single conditions.

Further, the optimal ‘cue weights’ predict subjects’ response location in double conditions from their response location in single cue conditions. This cue integration prediction can be explicitly tested when putting landmark cues into conflict by rotating them around the location of the last target. Doing so creates two response locations indicating exclusive use of either landmark or self-motion cues. Proximity of subjects response 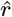 to either of those location, e.g. 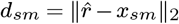. Where the self-motion consistent location *x_sm_* is either the true target’s location, e.g. (0,0), or when accounting for a particular subjects response bias by using the mean location of the self-motion consistent location [39], i.e., 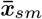. For the landmark-consistent location, the rotated normalized target location is used. It can further be corrected for subjects’ intrinsic response bias by subtracting the subjects’ mean response 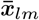 in landmark condition [39]. Given both proximities, i.e., *d_sm_* and *d_lm_*, to either of the single cue consistent location, we calculated relative response proximity to the landmark-consistent location as follows:

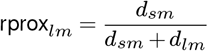

Response proximity to the landmark-consistent location is always between zero and one. Zero indicates subjects responding at the self-motion consistent location, whereas one indicates subjects responding at the landmark-consistent location. Relative response proximity can, therefore, also be interpreted as an empirical ‘cue weight’ subjects place on either cue, given homing responses in conflict conditions [39].

#### Ideal Observer Analysis (Heading Direction)

Zhao 2015 [38] performed cue integration analysis based on heading direction rather than euclidean distances. Heading errors were fitted to von-mises distributions for each cue condition separately (self-motion: 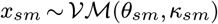; landmark: 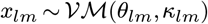; combined: 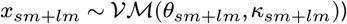. Predictions of mean heading direction and heading variability in combined cue and conflict conditions are predicted based on heading variability and heading directions from single cue conditions as follows:

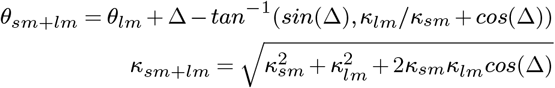

where *θ* refers to the mean homing direction and *κ* to the concentration. In conflict conditions, Δ refers to the angle of the landmark shift applied. Circular standard deviations are computed based on concentration parameters *κ* according to:

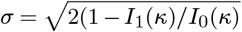

where I(x) refers to the Bessel function of the first kind.

### DYNAMIC BAYESIAN ACTOR MODEL

#### State Space Representations

The agents state **x_t_** consists of its pose (*x, y, θ*), including its position and heading direction, as well as the global (*x,y*)-coordinates of the landmarks (and target locations) in an allocentric frame of reference. The complete state thus can be described as **x** = (**x**_pose_, **x**_map_) where 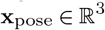 and 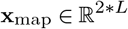:

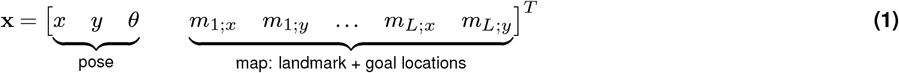

where L is the number of landmark and goal locations in the environment. The current state of the world at timestep t is expressed as **x_t_**.

#### Motion model and motor variability

The dynamic actor model changes its position and heading direction in space by sequentially performing motor actions. For the non-linear dynamics **x**_**t**+1_ = *f* (**x**_t_, **u_t_**) we assume a unicycle motion model with control inputs **u_t_** = (*v_t_, w_t_*). Prior research has shown that this non-holonomic motion model applies to human data for goal-oriented locomotion [12, 64, 113]. Here, we only model the discrete first-order dynamics with direct inputs of linear velocity *v_t_* and angular velocity *w_t_* for a discrete timestep dt:

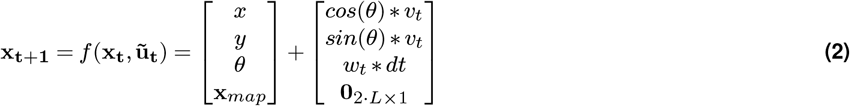

Further, we assume that these inputs *u_t_* = (*v_t_, w_t_*) are subject to signal-dependent noise *α* with noise parameters (*α*_1_, *α*_2_, *α*_3_, *α*_4_):

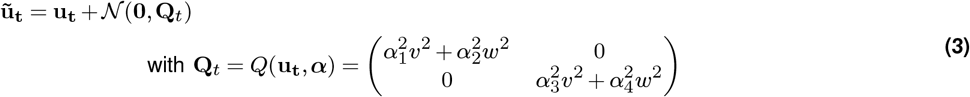

#### Observation model and observation variability

At each timestep, the agent makes noisy egocentric observations **z_t_**, i.e., relative distance and bearing [30], of landmarks and objects in the environment. When the agent encounters landmarks or objects for the first time, observations are converted to allocentric coordinates and then stored within the internal representation [99]. The non-linear observation model **z_t_** = *h*(**x_t_**) + **r_t_** consists of observations with additive Gaussian noise. Once in the field of view of the agent, an observation of a landmark l consists of its distance and direction relative to the agent, which can be expressed as:

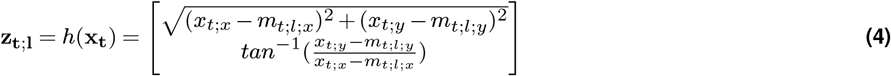

We assume that correspondence between a measurement and landmark l is unambiguously known. The agent observes Landmarks or targets if they fall inside the field of view of the agent. Thus the complete environment is not observable to the agent at once. Further, observations are subject to state-dependent gaussian noise with noise parameters ***ε*** = (*σ_r_min__, σ_r_max__, σ_ψ_min__, σ_ψ_max__*):

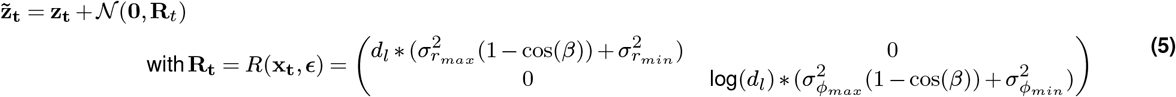

where *d_l_* refers to the distance and *β_l_* refers to the angle between the agent’s gaze direction and the vector between the agent and the landmark or target. This dependence on angle leads to observation in the visual periphery being noisier compared to observations made in the center of the field of view [48]. The dependence on distance makes observations noisier with increasing distance [8].

#### Belief Space Representation

As the true state of the world can only be perceived indirectly, the agent only has access to the belief state *b_t_* = (**μ_t_**, **Σ_t_**) capturing the mean location of their location, heading direction and the locations of objects, landmarks and their associated uncertainty. This extends the state-space formulation from Equation 1 by an explicit representation of spatial uncertainty over the agent’s position, heading, landmark, and goal locations.

The belief **b_t_** at time-step t describes this as follows (t dropped for notational convenience):

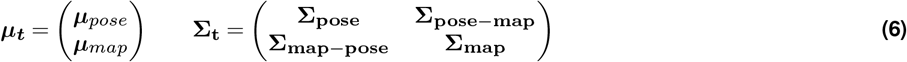

with the following submatrices:

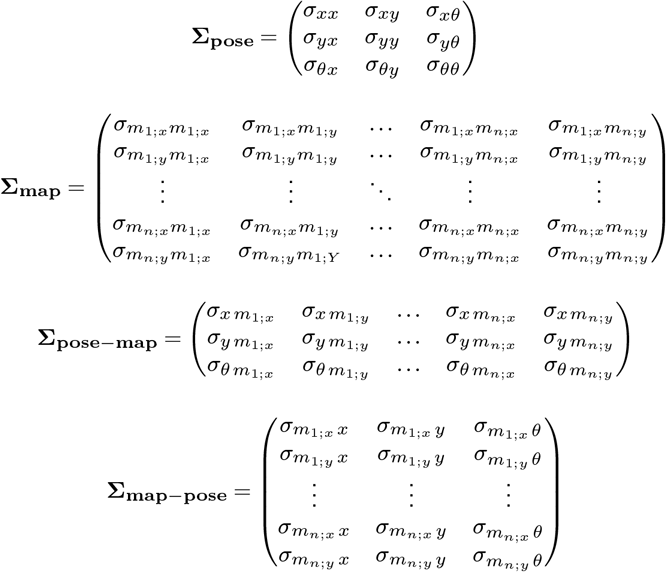

Having access to the full belief state, rather than a single point estimate of self and landmark locations allows for the model to keep track of the growing positional uncertainty and resolve it later by actively reorienting and thereby combining noisy observations of landmarks with prior expectations about the position.

#### Bayesian Filtering: Belief Dynamics & State Estimation

The framework of Bayesian filtering provides a principled manner in which the belief state is updated as a consequence of perception and action [26, 47, 57]. Central to these computations are the two steps of prediction and correction. The prediction step uses a probabilistic motion model of the form *p*(*x_t_*|*x*_*t*−1_, *u_t_*) and increases positional uncertainty. The correction step uses an observation model of the form *p*(*z_t_*|*x_t_*), which incorporates noisy landmark observations from the environment with prior uncertain expectations about the landmarks’ locations stored in the internal representation. Both steps can be performed by recursively applying the probabilistic Bayes Filter:

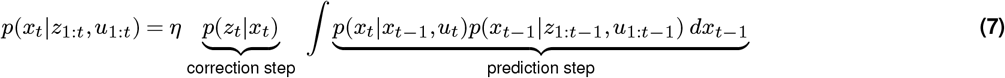

Choosing functions the non-linear dynamics and observation functions f and h such that they are differentiable allows for local linearization around the current state and application of an Extended Kalman Filter [57, 114]. This allows for a closed-form belief propagation of the form (***μ_t_***, **Σ_t_**, **u_t_**, **z_t_**) → (***μ***_*t*+1_, **∑**_**t**+1_).

The equations for the **prediction step** then become:

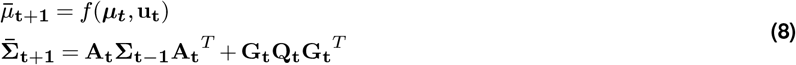

where

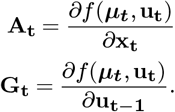

Representation uncertainty **V_t_** gets added to landmarks *m_L_* ≔ (*m_L;x_, m_L;y_*) at each timestep

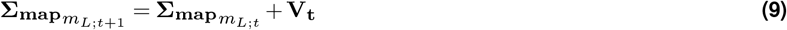

where

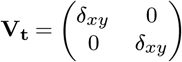

The equations for the **correction step** then become:

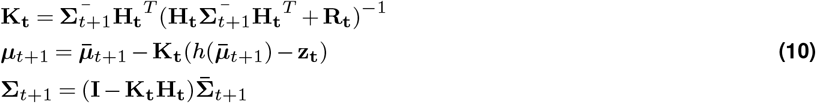

where

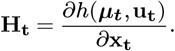

#### Representation Variability

We assumed that the agent’s internal representation is noisy, such that precise memory of objects and landmark location decays over time [115]. Landmarks *m_L;t_* ≔ (*m_L;x_, m_L;y_*) follow a 2D random walk [116] according to:

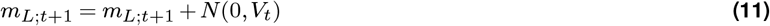

#### Belief Space Planning

The agent plans motor actions based on its current belief about its pose and the location of landmarks within the environment (***μ_t_***, **Σ_t_**) by propagating it according to the belief dynamics while minimizing cost function J [53, 114, 117]. The planning step results in a sequence of states *μ*_0:*T*_ and actions *u*_0:*T*–1_ and can be written as a constrained non-linear trajectory optimization problem of the form:

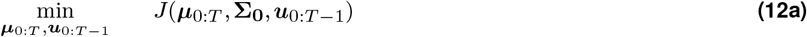

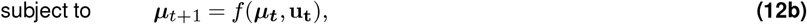

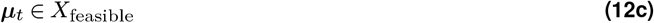

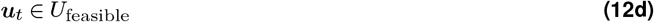

where T refers to the planning horizon and J to the cost function. Feasible values for the optimization are those which satisfy the following lower and upper bounds:

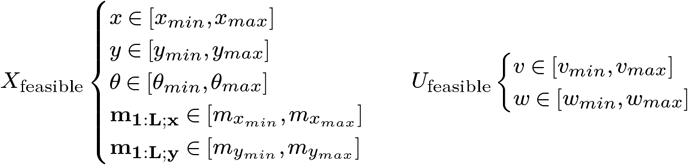

We only include the mean states ***μ_t_*** as well as the control inputs **u_t_** in our optimization problem formulation. Excluding the covariance from the optimization variables reduces the complexity of the planning problem, thus speeding up runtime [117]. This allows for effective re-planning at every timestep, which is necessary to incorporate closed-loop feedback into the system via model predictive control. Under this set of assumptions, the problem of belief space planning becomes open-loop and deterministic. In order to deal with future observations, yet unknown to the agent during planning, we assume that any stochasticity is due to gaussian noise, so the agent uses maximum likelihood observations during planning, i.e., **z_t_** = *h*(***μ_t_***) [53].

The restricted field-of-view of the agent leads to partial observability of the environment, which needs to be considered by the agent during planning to select appropriate actions that reduce uncertainty about task-relevant quantities. In order to include landmarks currently not visible in planning future actions, we change Equation 10 for the planning problem. We include Δ_*t*+1_, a diagonal matrix comprising of a vector *δ*_*t*+1_, which indicates which landmarks are currently observable to the agent and which are not (maximum likelihood observations). Rather than using a strict indicator function to determine the visibility of a landmark, its visibility is approximated in an iterative fashion using a sigmoid function, i.e., 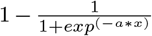, for decreasing values for a, together with signed distances x to each object and landmark relative to the agent’s current belief about the position and heading. This changes the computation of **K_t_** for the planning belief dynamics in the update step as follows:

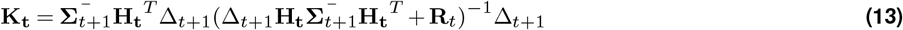

#### Cost Function

Which trajectory the agent plans and executes is strongly influenced by the choice of the cost-function. A cost function for navigation should entail reaching a target in space (a), reducing the uncertainty about self-location (b), and the location of goals and landmarks (c) while penalizing control effort (d). This leads to several different terms within the cost function J. Given a set of feasible states and controls ***μ_t_*** and an internal belief about the location of the goal **g_t_** = (*m_μ_l; x, t__, m_μ_l; y, t__*), the objective function *J* can be computed by propagating the initial belief state **b**_0_ = (***μ***_0_, ***∑***_0_) with the sequence of controls *u*_0:*T*_ using Equations 8 and 10:

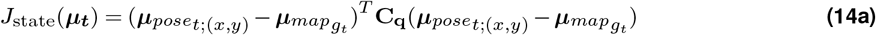

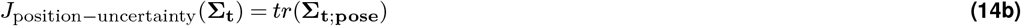

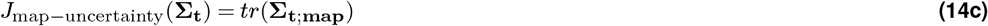

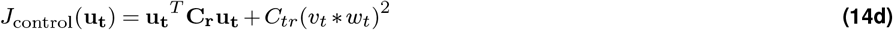

This leads to the following weighted cost function for the planning problem:

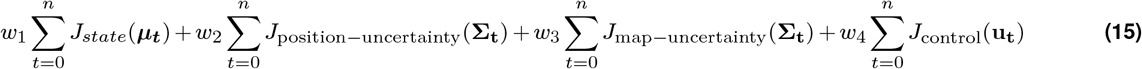

with weights **w** = (*w*_1_, *w*_2_, *w*_3_, *w*_4_). This set of weights captures the general trade-off between sensing, i.e., remaining oriented by reducing uncertainty about agent position and landmark position, and acting, i.e., reaching a goal. Once the trajectory is planned for T steps (planning horizon), the agent executes the first N steps of the current plan (control horizon).

#### Reorientation

Due to the nature of the task and noise in the motor system, the agent’s internal model and the true state of the world could become so divergent that the agent’s planning procedure may get stuck. For example, suppose the agent is completely disoriented. In that case, e.g., it expects to see landmarks based on the belief about heading direction and internal representation about landmark location but does not observe any landmarks. In this case, the agent will keep turning in the direction it believes landmarks are until it brings them into the field of view. The observation is then integrated with the internal model via Equation 10, providing reorientation so that the agent can further plan and act accordingly to its beliefs about the world.

### METHOD DETAILS

#### Simulation of Homing Task Conditions

For the homing task, the simulation of the agent navigating the outbound path was the same for all conditions. Depending on the state of the task, different target locations were shown. Targets were either visible to the agent simultaneously from the beginning of the trial or appeared in succession during the unfolding of the triangle completion task. Both the location of landmarks and targets were initialized at the location where the virtual agent first encountered them. Each trial was simulated identically and independently, i.e., with no trial-by-trial learning. Before homing, the agent had to wait for 8-20s, depending on the experiment. We modeled the different cue conditions during the homing response by manipulating the reliability or availability of the different cues. In self-motion conditions, we removed the availability of landmark cues during the homing path. In landmark conditions, we disoriented the agent by simulating a rotating swivel chair. For this, we first applied a slight positional offset, which we extracted from trajectory data (if available) or otherwise assumed a gaussian distribution similar in magnitude. After this, we simulated the agent’s disorientation by rotating them in their location using the turning speeds of the original studies. Since disorientation has also been shown to affect subjects’ internal representation of landmark locations [118], we increased the agents’ uncertainty about landmark locations by applying a small amount of process noise (see Equation 9). In combined conditions, we had the agent wait for n seconds before homing. Similarly, the agent remained still in conflict conditions for n seconds, and landmarks were rotated x degrees around the location of the third post before homing.

#### Model Parameters & Simulations

Values of noise parameters were constrained by findings from previous experiments in the literature on locomotion [12] and perception [8] and further manually fitted to trajectory and endpoint data for all experiments simultaneously (see supplementary material for model parameter table), placing strong constraint on the model parameters, being able to generalize across multiple studies and experimental manipulations. For each of the experiments, we obtained the number of trials for each of the four conditions, the subjects starting position and sequence of goal locations, and other relevant differences between environments and task procedures (see Table 1). The optimal control formulation (see Equation 12) was implemented in the CasADI framework [119], which allows for the specification of complex cost-functions and automatic differentiation. Planning is achieved by repeatedly solving a non-linear program (see Equation 12) under the set of constraints Equation 3) using IPOPT and the HSL solvers (academic license). Trials were executed in parallel on a compute cluster using Ray Tune [120], which allowed for efficient scheduling and logging of individual trial results.

#### Model Comparison

To further understand the complex interaction of the three sources of sensorimotor noise, we compared endpoint variability from different versions of the dynamic actor model to empirical data by selectively excluding particular sources of variability. These models either considered variability in perception, representation and action jointly (full model), separately (perceptual variability only; motor variability only; representation variability only), without one of the sources (perceptual variability & representation variability; representation & motor variability; motor & perceptual variability), or not at all (zero model). Each of the eight different models was simulated for each experiment 10 times. For each endpoint distribution within the four conditions, we computed the energy distance [121] between the observed distributions in the experiment and the simulated endpoint distributions from each model. This allowed us to assess how well each of the different models can account for the endpoint variability observed in the respective experiments, and how much and in what manner the three sources of variability contribute to the observed endpoint variability.

### QUANTIFICATION AND STATISTICAL ANALYSIS

#### Data Preprocessing & Analysis

All endpoint data, i.e., empirical data from the respective studies and data from model simulations, were reanalyzed following the procedures used in prior work [35, 38, 39]. We normalized homing responses for each homing location by centering them within a common coordinate system, with either the actual homing direction as the north-direction [38] or centered around the true targets location[35, 39]. Outliers were removed based on distances to the mean location within each condition by excluding those trials which exceeded the third quartile by three times the interquartile range [35, 39]. For data of Chen 2017 [39] and Zhao 2015 [38], we also removed biases in the four individual targets locations before pooling them together so as not to increase response variability artificially [39]. After pooling data from all four target locations, we computed the response variability for each (simulated) subject in each of the four conditions. We did so either as the standard deviation of the euclidean distances (or heading directions) between the response locations for each trial to the mean response location (or heading) across all trials. We used computed response variances for the two single cue conditions (self-motion and landmark) to predict response variance for the combined cue condition according to the perceptual cue integration model for euclidean distance [35, 39] or heading direction [38]. We considered the integration of cues statistically optimal when the reduction in response variability in the combined cue conditions can be predicted from response variability in single cue conditions. Further, we used mean responses in conflict conditions to predict relative response proximity to the landmark or self-motion-defined target locations. We compared them with optimal predictions based on response variability. All data were analyzed at the single subject level if not indicated otherwise.

#### Statistical Tests

We tested endpoint distributions for the different cue conditions for multivariate normality, but this assumption was violated for most endpoint distributions. Therefore, we resorted to using energy statistics [121] from the python package dcor, which allows for quantifying the degree of similarity (via energy distance) and testing for equality (via homogeneity energy test) of random vectors sampled from arbitrary multi-dimensional non-parametric distributions. We compared response variability between empirical and simulated data based on mixed ANOVAs using the python package pingouin [122]. For comparing response variability between model and empirical data, we performed two-way repeated measures ANOVA on response variability with type (model simulation vs. empirical data) and cue condition (self-motion, landmark, combined, conflict) as independent variables. We performed Posthoc tests using pairwise t-tests. We considered results significant if p < 0.05. Multiple comparisons were corrected using Bonferroni corrections only if indicated in the main text.

## Supplementary Information

**Figure S1.**
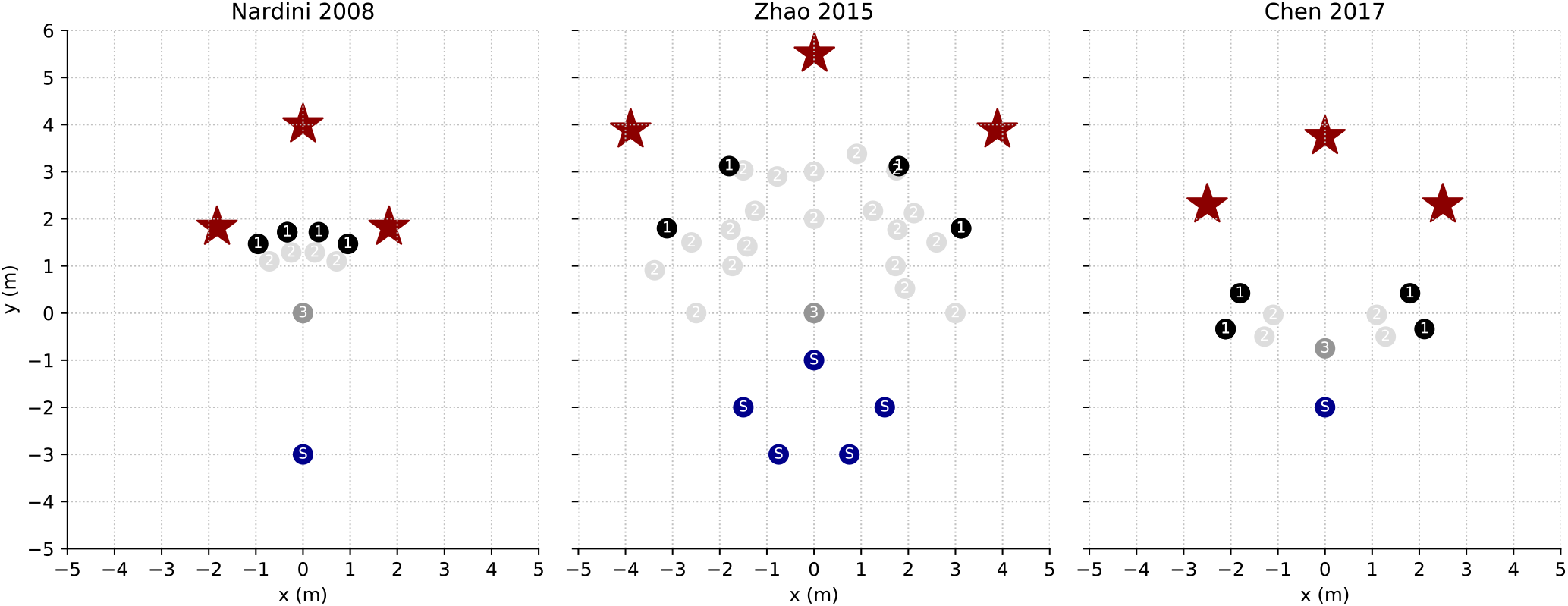
Environmental layouts in [1–3] with location of starting position, goal posts and landmarks. Different combination of goal locations lead to different three legged outbound paths.

**Figure S2.**
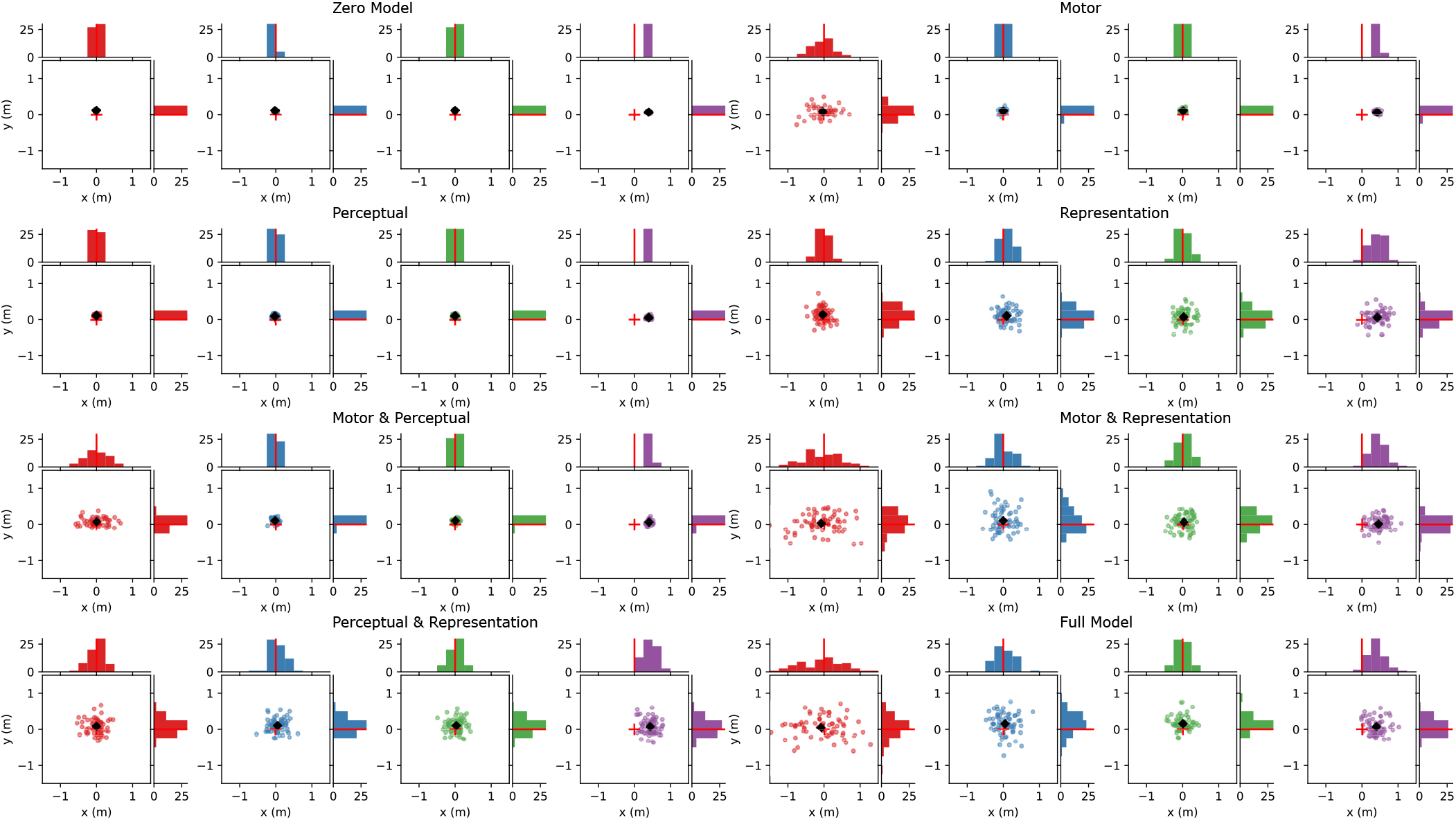
Nardini 2008 Model Comparison: Endpoint distributions.

**Figure S3.**
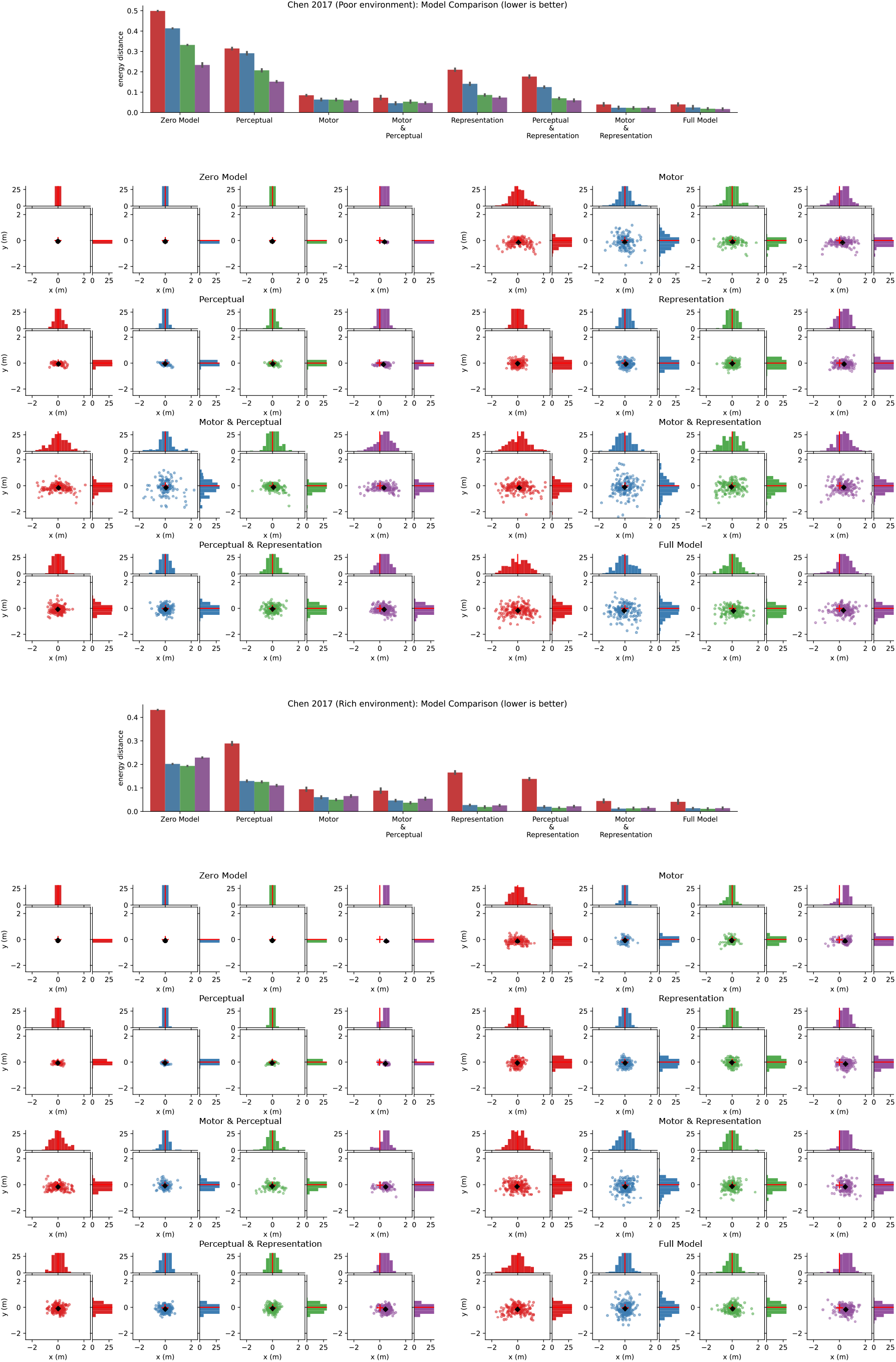
Model Comparison Endpoint Distributions Poor & Rich Environments (Chen 2017)

**Figure S4.**
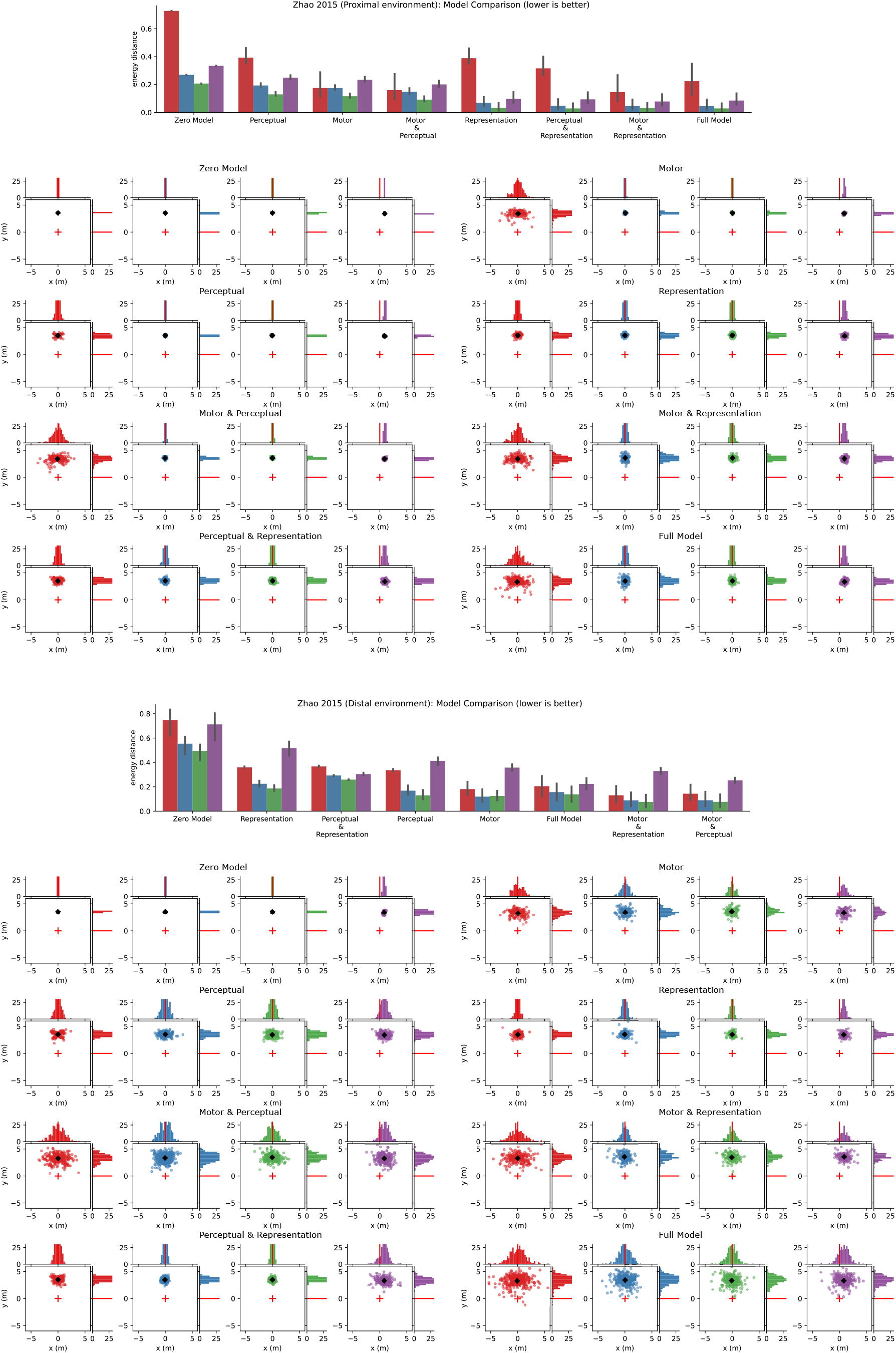
Model Comparison Endpoint Distributions Proximal & Distal Environments (Zhao 2015)

**Figure S5.**
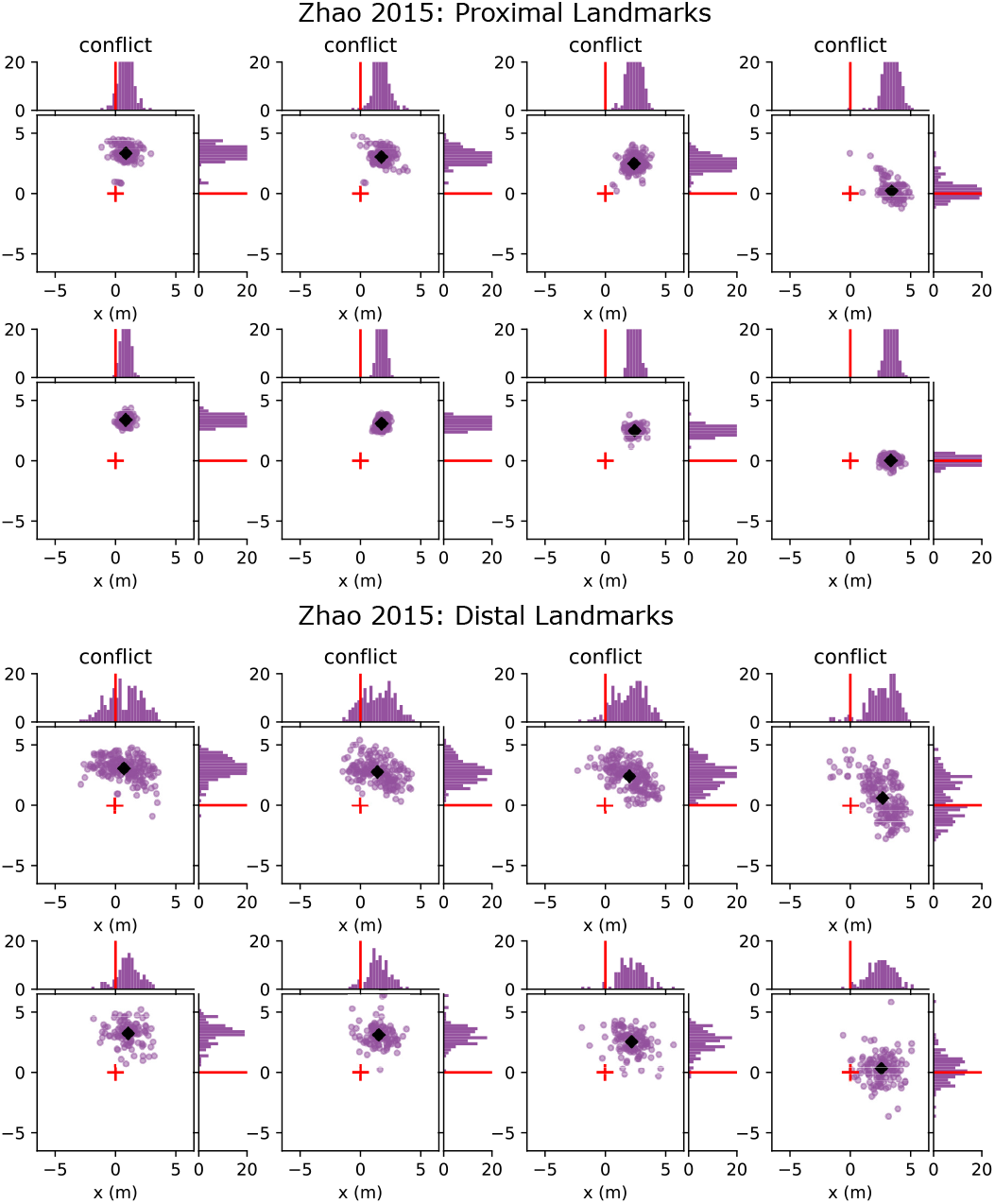
Zhao 2015: Conflict Conditions 15-90°.

